# Systematic transcriptomic analysis and temporal modelling of the senescent human fibroblast

**DOI:** 10.1101/2022.08.16.504144

**Authors:** R-L. Scanlan, L. Pease, H. O’Keefe, A. Martinez-Guimerá, D Shanley, J Wordsworth

## Abstract

Cellular senescence is a diverse phenotype characterised by permanent cell cycle arrest and an inflammatory senescence associated secretory phenotype (SASP). Typically, senescent cells are removed by the immune system. This process becomes dysregulated with age and senescent cells accumulate leading to chronic inflammatory signalling. Identifying senescent cells is challenging due to the heterogeneity of senescence, and senotherapy often requires a combinatorial approach. Here we have taken an integrative approach to investigate senescence development at the transcriptomic and protein level. We systematically collected 119 transcriptomic datasets related to human fibroblasts, forming an online database describing the relevant study variables which users can filter to select variables and genes of interest. Our own analysis of the database identified 28 genes significantly up- or downregulated across four senescence types (DNA-damage induced senescence (DDIS), oncogene-induced senescence (OIS), replicative senescence, and bystander induced senescence); 14 genes consistently downregulated, 10 genes consistently upregulated and 4 genes regulation dependent on senescence type. We also found gene expression patterns of conventional senescence markers were highly specific and reliable for different senescence inducers, cell lines, and timepoints. Conclusions of existing studies based on single datasets were supported such as differences in p53 and inflammatory signals between DDIS and OIS. However, contrary to some early observations, both p16 and p21 mRNA levels appeared to rise quickly, depending on senescence type, and persist for at least 8-11 days. Additionally, little evidence was found to support an initial TGF-β-centric SASP. To support our transcriptomic analysis, we computationally modelled temporal protein changes of core senescence proteins in DDIS and OIS, as well as performed knockdown interventions. We conclude that while universal biomarkers of senescence are difficult to identify, conventional senescence markers follow predictable profiles and construction of a workflow for studying senescence could lead to more reproducible data and understanding of senescence heterogeneity.

## Introduction

Multiple studies now suggest that the accumulation of senescent cells is causal in ageing (Childs et al., 2015; Mylonas & O’Loghlen, 2022; van Deursen, 2014; Wlaschek et al., 2021), and their ablation extends healthspan and mean lifespan in rodents (Baker et al., 2016; Baker et al., 2011). Novel senolytic and senostatic drugs are in development (Kim & Kim, 2019; Niedernhofer & Robbins, 2018), with some drugs in clinical trials (Hickson et al., 2019; Justice et al., 2019) which might shortly lead to treatments capable of improving healthspan and extending lifespan in humans. However, the exact nature of senescent cells is often difficult to define, with multiple studies indicating that the most common biomarkers of senescence show different profiles across cell lines, types of senescence inducer, and the timepoint after the initial stimulus (Avelar et al., 2020; Basisty et al., 2020; Casella et al., 2019; Hernandez-Segura et al., 2017; Neri et al., 2021). This makes targeting senescent cells difficult, often requiring combinatorial approaches (Nayeri Rad et al., 2022; Saccon et al., 2021; Xu et al., 2018; Zhu et al., 2015). Although combination therapies can be effective, they also have potential to impact additional molecular networks and their off-target effects can be unpredictable.

To gain a comprehensive understanding of cellular senescence, an integrative, systematic, multi-omic approach is needed. Consequently, we have taken an integrative approach and conducted a transcriptomic systematic review of cellular senescence in human fibroblasts alongside development of a qualitative protein computational model which simulates DNA damage induced senescence (DDIS) and oncogene induced senescence (OIS). We have systematically analysed all transcriptomic data for senescent fibroblasts, meeting pre-specified inclusion criteria, and produced an online database that allows public analysis of the results. Firstly, changes across four types of senescence were compared before analysing temporal changes in key senescence biomarkers in DDIS and OIS, comparing our results to existing single dataset studies. We then examined the literature to build a qualitative protein level model of cellular senescence, using the transcriptomic profiles observed here to aid in the development of the model. For validation of the model, knockdown (KD) interventions of p53 and RelA were performed and compared to data.

This integrated approach has allowed us to address universal biomarkers for senescence, whether findings from individual studies are consistent across the total data, and the mechanisms by which the network of interactants might induce the senescence response. Our results indicated differences in DDIS and OIS for expression of several of the key senescence genes, but profiles were largely consistent with results from individual studies. One exception was the TGF-β-centric SASP identified by Hoare et al. (2016), which does not appear to be replicated in the total data at the transcript level.

## Methods

### Systematic review protocol

Three independent systematic searches were conducted and updated to identify all transcriptomic data meeting our inclusion criteria for cellular senescence in human fibroblasts publicly available by 05 October 2023. Datasets were included if they met the following inclusion criteria:

- Unbiased transcriptomic datasets for senescent human fibroblasts. Where senescence was defined exclusively by permanent cell cycle arrest induced by a stimulus in a cell type that would otherwise be proliferating.
- RNA-seq or microarray datasets stored on Gene Expression Omnibus (GEO) (Edgar et al., 2002) or Array Express (Parkinson et al., 2007) by the deadline date of 05 October 2023.
- Data had at least two repeats for all conditions included.

As some datasets meeting the inclusion criteria could not be analysed by the methods described below, they were further excluded if they met the following exclusion criteria:

- Exclusively microRNA or long non-coding RNA datasets.
- Performed at the single cell level.
- Two colour or custom microarrays.
- Data could not be downloaded from GEO or Array Express, nor provided by contact with the corresponding author.

Search terms were developed to include all relevant MeSH terms and text terms that might identify datasets meeting the inclusion criteria. Initial terms were used in combinations on PubMed PubReMiner (Slater, 2014) to identify additional search terms. The search terms selected for GEO and search results of the initial and updated searches are shown in **Table S1** used in the Advanced Search tool to combine individual searches.

For the smaller Array Express database we searched for ‘Ageing’ OR ‘Aging’ and then manually filtered the results.

As described, manual exclusion was done for both databases in three independent searches firstly on 10 August 2020. At this time the results were compared to an initial non-systematic search of both databases as well as a PubMed search for studies including transcriptomic data, producing a control dataset that ensured the systematic search identified all the studies in the preliminary search. The initial systematic search was then followed by two updated searches, on 06 July 2022 and 05 October 2023. All three searches were done by the same two individuals for two independent searches per search date. After each search, the results were compared to those of the other individual and previous searches to ensure that no studies were missed.

A total of 5,095 studies were identified up to 10 August 2020. Duplicates were removed and remaining studies were reviewed manually against specific inclusion and exclusion criteria. There was an initial total of 82 studies identified. The updated searches on 06 July 2022 and 05 October 2023 following the same search criteria identified a further 16 studies and 21 studies respectively; resulting in a total of 119 studies included in the systematic database.

### Database Creation

For each study, a comparison matrix was constructed in Microsoft Excel listing all data of interest that could then be combined into a single searchable database. If the data of interest were not available on GEO or Array Express and the datasets had accompanying publications, we checked the papers for any missing data. Key data such as senescence type and cell line were available for all datasets; however, in some cases, the timepoint of senescence induction was not stated in the paper or online databases. As this was a key part of constructing a senescence profile, we then contacted the corresponding author, but we did not do so for any other missing categories.

### Data preparation and analysis

RNA-seq data was downloaded as fastq files from GEO or Array Express. Each file underwent quality check using the fastqcr R package (de Sena Brandine & Smith, 2019) (in R version 3.6.3) and files were compared using the MultiQC BASH command (Ewels et al., 2016). Adapter trimming and removal of low quality read ends was carried out using the Cutadapt tool (Martin, 2011). Once fastq files passed fastqc, or were excluded, they were converted by mapping-based quantification to quant.sf files using Salmon (version 1.1.0) (Patro et al., 2017). The --gcBias --seqBias and -- validateMappings options were used to remove additional biases.

For microarrays, series matrix files for selected studies were downloaded from GEOquery and loaded into R using GEOquery (Davis & Meltzer, 2007), converted to esets and labelled with normalisation and processing information provided with the files. Array Express raw data sets were downloaded using ArrayExpress and RMA normalised using affy (Gautier et al., 2004).

Quant.sf files and microarray data underwent differential expression analysis using the R limma package (Ritchie et al., 2015). Data were normalised by cpm or voom commands depending on variance, and plotDensities was used to compare sample curves. Samples with irregular curves not consistent with the rest of the data were removed from further analysis. Log fold change (LogFC) and p values were calculated for each comparison defined in the comparison matrix using the eBayes function and combined into a single database for all studies (available on website, see below).

During initial analyses it was noted that the LogFC of some genes were clear outliers. To prevent outliers skewing the analyses, the interquartile range (IQR) was calculated for each gene per timegroup. Outliers were identified as anything less than Q1 (25^th^ percentile) - 1.5*IQR or more than Q3 (75^th^ percentile) + 1.5*IQR. Outliers were removed prior to all analyses presented in this paper.

For some analyses p values were inverted (p_i_ value) by the formula in equation 1. This created a scale that put p values for significant upregulation at the opposite end to p values for significant downregulation, with non-significant values in the middle.

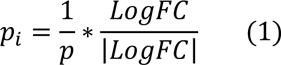

Gene set enrichment analysis was carried out using the GSEA command from the ClusterProfiler library (Wu et al., 2021), using the GSEA index h.all.v7.0.symbols.gmt.

We employed the Wilcoxon signed-rank test (which comes in the base R software) for paired sample statistical analysis (Bauer, 1972).

### Online database creation

We transformed the database into a Power BI report. This allows users to access various clusters of the data in easily readable visuals. Data clusters are accessible by button selection and users can sift the data through 13 different filters provided in the report. We used Power BI basic functions and DAX programming language to build the report; specifically, DAX was used to create measures which control the filtering selections. The Power BI report is embedded via an iframe in a Newcastle University research website, available at: https://research.ncl.ac.uk/cellularsenescence. The website holds subsidiary information on the report, project and research team.

### Computational model development

Computational model networks were developed for the purpose of qualitatively simulating temporal protein expression changes in DDIS and OIS. A protein network of reactions and interactions in cellular senescence was designed in CellDesigner (Funahashi, 2008), informed from extensive literature searching of temporal protein profiles in human fibroblasts (**Table S2**). When there was no available published protein data, transcriptomic data from our systematic database was used as the best available proxy for temporal protein behaviour. To model this protein network, we used Tellurium (Choi et al., 2018) in a Python 3.6 environment. KDs (p53 and RelA) were also introduced into all model networks to investigate whether senescent KD phenotypes could be successfully recapitulated.

Criteria for the model to meet were devised from commonly accepted senescence phenotypes in the literature as well as from the systematic analysis. Models were built and developed in stages; introducing more proteins, reactions, and therefore complexity, to create a model which would meet all devised criteria and behave sensibly and in accordance with the literature and systematic analysis. KD phenotype criteria were devised based on the transcriptomic systematic analysis. Meeting of these criteria is how we measured if a model network could successfully simulate cellular senescence.

To induce senescence specific inputs were utilised (**Table 1**). The inputs were selected and devised based on knowledge from published literature. All expression and time units in the simulations are arbitrary (AU).

**Table 1.** Inputs for simulating different cellular states. *Induced once the model had reached equilibrium.

| Inputs | Cellular states simulated |  |
| --- | --- | --- |
|  | DDIS | OIS |
| RAS | 0.5 | 5* |
| DDR | 5* | 0 |
| kDDRf | 14* | 0.01 |
| kRASf | 0.3* | 1* |
| kDDRFRAS | 0.5 | 5.75* |

Additionally, an inducible event termed the Notch switch was introduced into the final model network. The Notch switch creates a dynamic change in Notch signalling activity. Activation of the Notch switch results in diminished Notch signalling through the downregulation of NICD levels. Similar to the Notch switch, KDs were simulated through an inducible event. For simulation of a p53 KD, p53 levels and formation of p53 was downregulated. Likewise, for RelA KD simulations NF-κB levels and formation of active NF-κB were downregulated. KDs were induced at the same time as senescence induction, a decision based on the methods described in the p53 and RelA KD studies in the transcriptomic database.

Dynamic sensitivity analysis was performed on model C, the final model developed (also referred to as the Senescence Induction Model from here onwards), in both DDIS and OIS conditions at four timepoints (pre-senescence, T20 (Time 20); post-senescence induction, T40; pre-Notch switch, T60; post-Notch switch, T80). We followed the X-method by (Yue et al., 2006) for calculating scaled sensitives to parameter perturbations as ‘events’ at specific simulation time points using SIMBIOLOGY, MATLAB. Results from dynamic sensitivity analysis were as expected, with all proteins appropriately sensitive to senescence inputs and the Notch switch event at the expected timepoints (**Figure S1**).

## Results and Discussion

The systematic search criteria outlined in the methods initially identified 5063 studies from GEO and 32 studies from Array Express. Of these, 26 were removed as duplicates leaving 5069 datasets for manual analysis. Of these, 82 were identified as meeting the inclusion criteria and not disqualified by the exclusion criteria. The update searches identified a further 36 studies, as shown in the PRISMA flowchart in **Figure 1**. A total of 119 studies, including 85 RNA-seq datasets and 34 microarray datasets, were identified.

**Figure 1.**
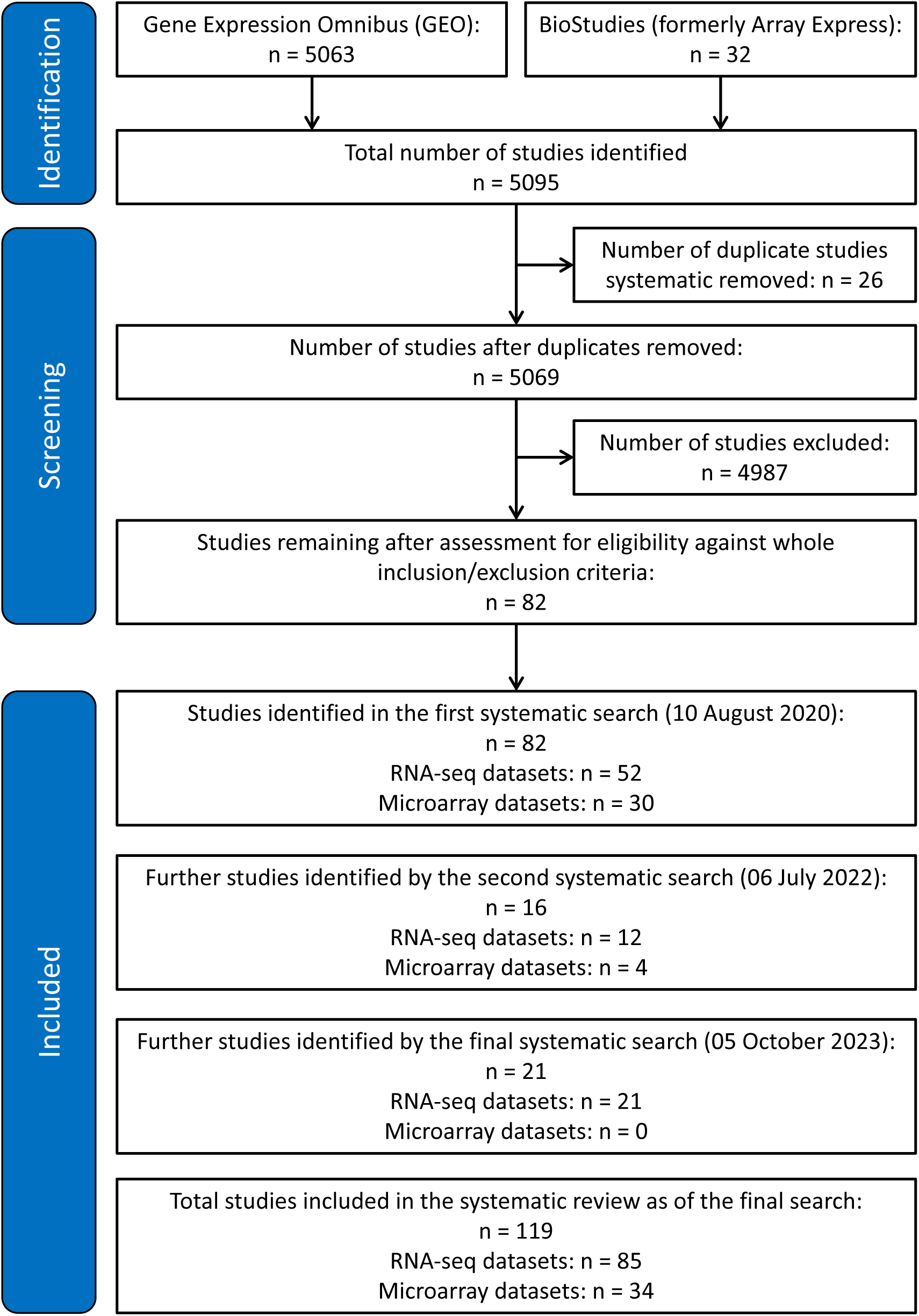
PRISMA flowchart showing identification and exclusion of studies.

From these 119 studies we made a total of 1069 comparisons, 220 of which were between senescent cells and proliferating controls without treatment or disease. The details of the studies included are shown in **Table 2**. The main categories, acronyms, and number of comparisons for each are shown in **Table S3**.

**Table 2.**
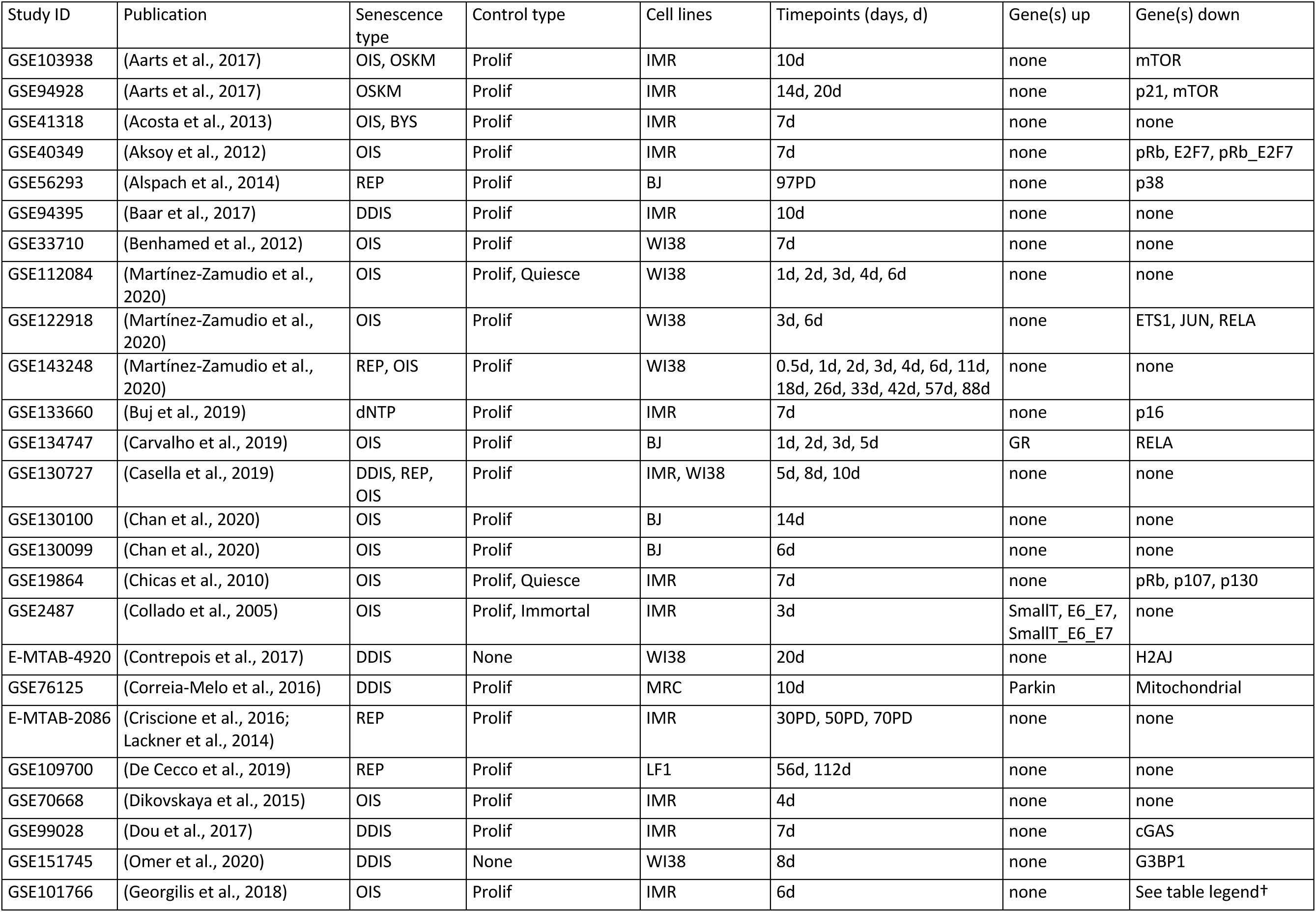
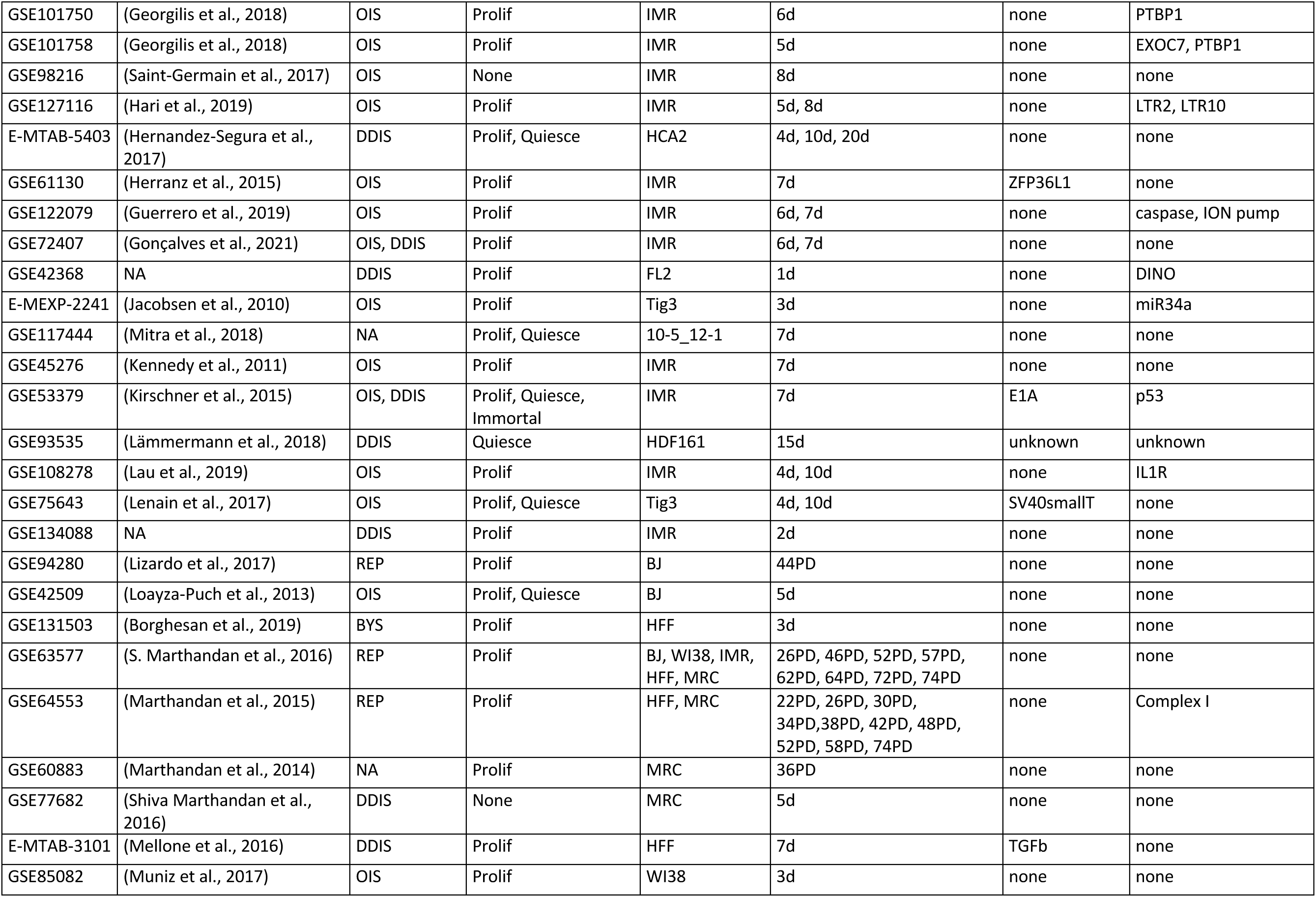
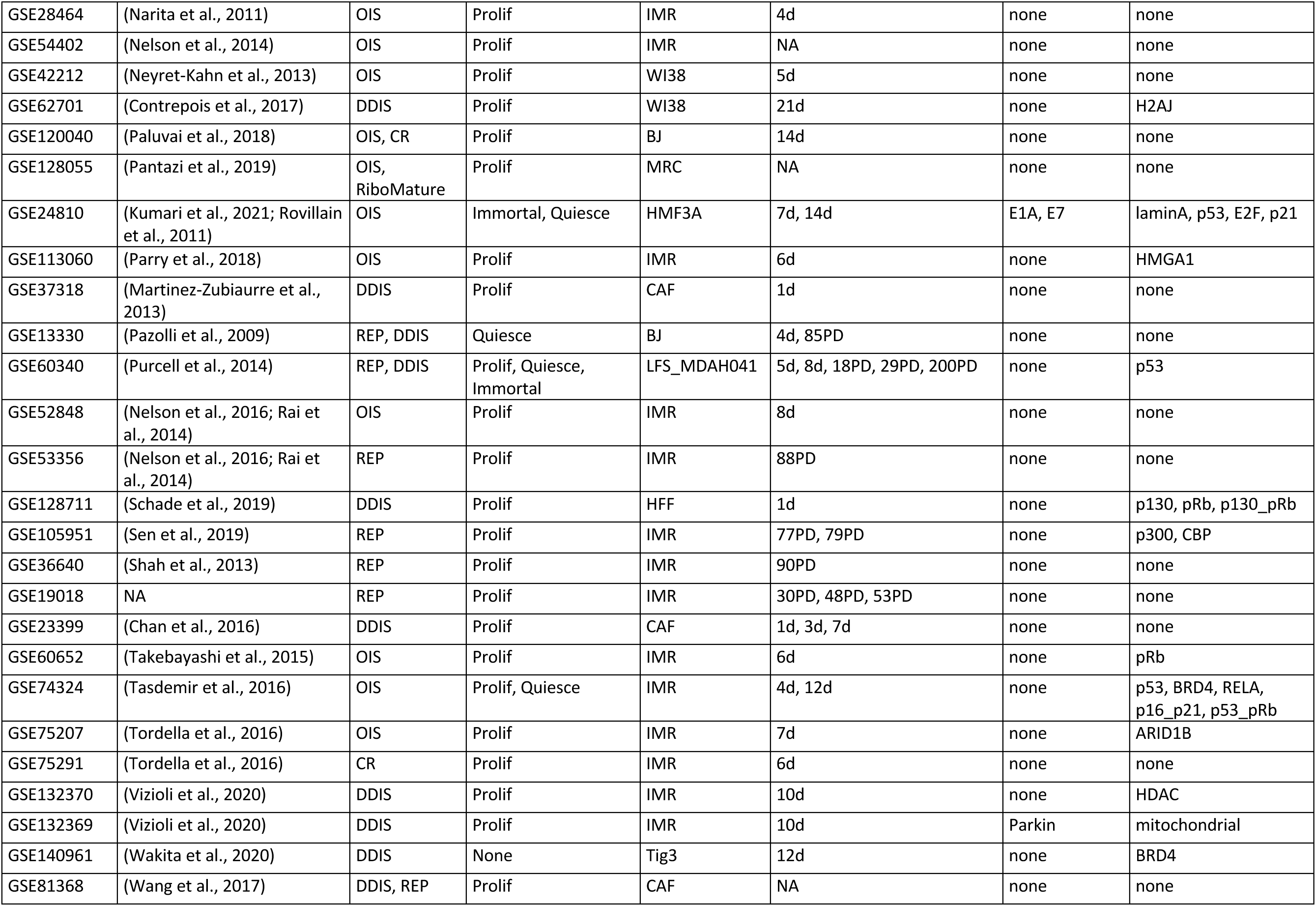
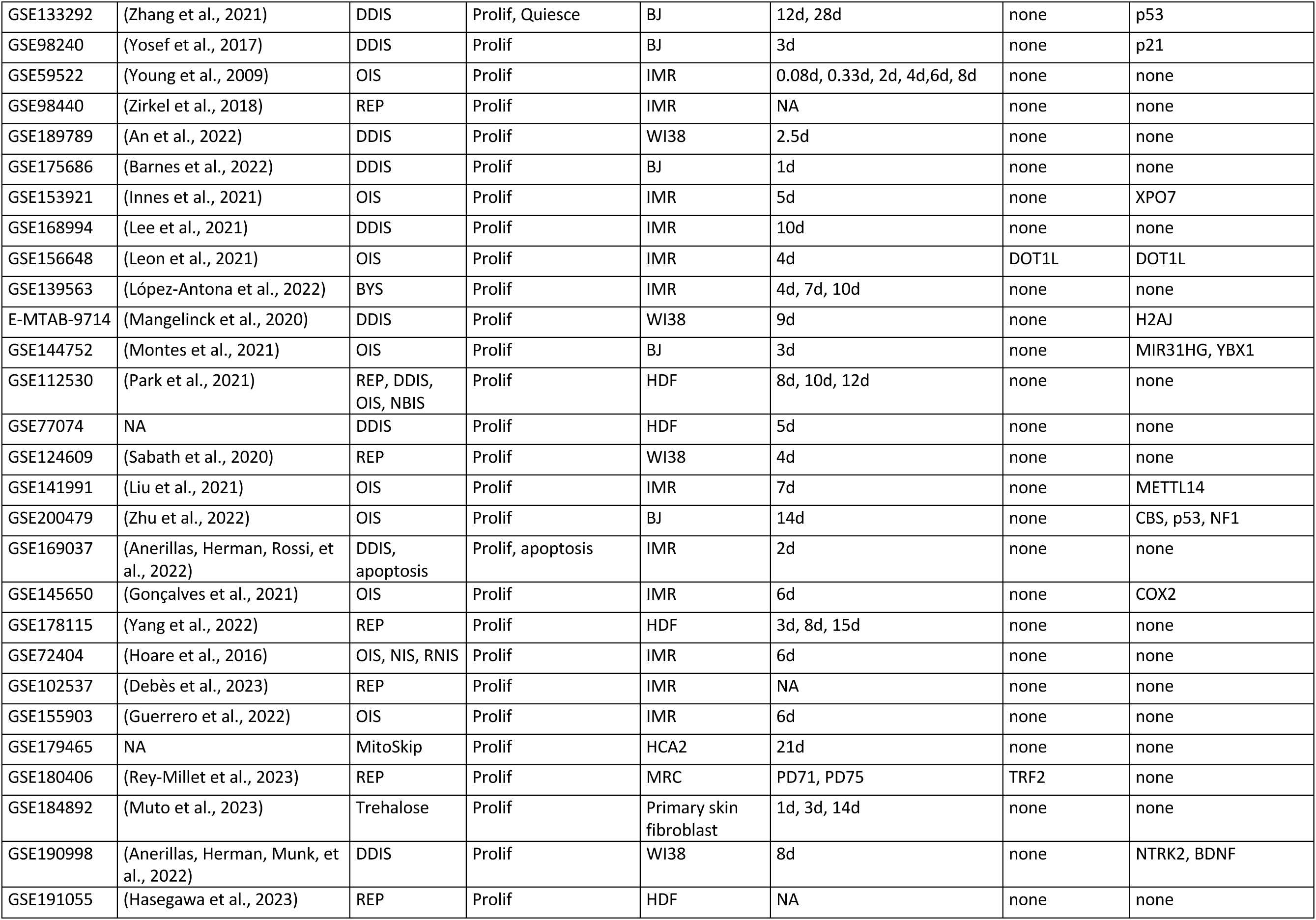
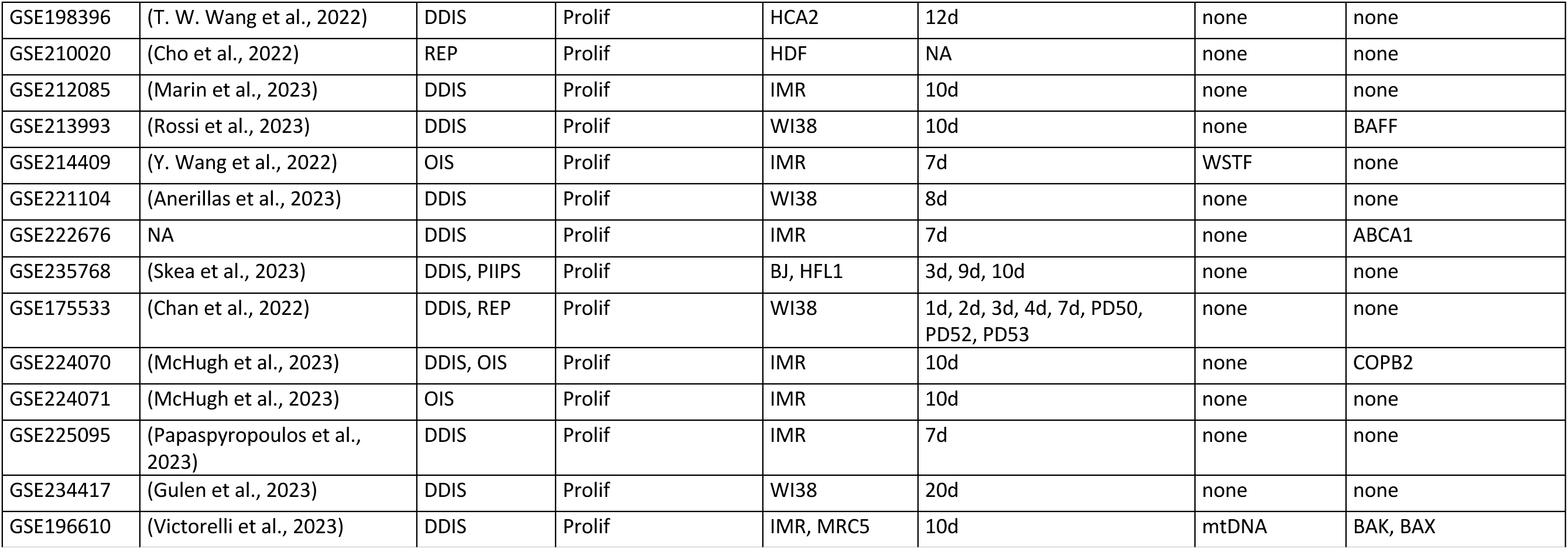
Study data for the 119 included studies. Prolif, proliferating cells; Quiesce, quiescent cells; Immortal, immortalised cells; PD, population doublings; CAF, cancer associated fibroblast; BYS, bystander induced senescence; DDIS, DNA damage induced senescence; OIS, oncogene induced senescence; REP, replicative senescence; CR, chromatin remodelling induced senescence; NBIS, nuclear breakdown induced senescence; NIS, Notch induced senescence; RNIS, Ras and Notch induced senescence; OSKM, senescence induced as a by-product of pluripotency induction via transcription factors Oct4, Sox2, Klf4 and c-Myc; dNTP, depletion of deoxyribonucleotide triphosphates; RiboMature, senescence induced through ribosomal disruption; MitoSkip, senescence induced via mitotic skipping; PIIPS, proteasome inhibition-induced premature senescence; Trehalose, senescence induced though high concentrations of trehalose. †CEBPb, ABCD4, AKR1C1, ALOX5, ASB15, BPIL1, BRD8, C20, CCL23, CTDSPL, DCAMKL3, DUSP11, EMR4, ERCC3, GPRC5D, HSPC182, IFNA17, IL15, IL17RE, ITCH, KCNA5, KCNQ4, LOC399818, LOC51136, MAP3K6, MCFP, NRG1, PEO1, PLCB1, PPP1CB, PROK2, PTBP1, PTPN14, RNF6, SHFM3, SKP1A, TMEM219, UBE2V2, p16, p38, p53, RELA.

Fourteen types of senescence induction were identified from the literature and included in the database: replicative senescence (REP) from telomere erosion (Bodnar et al., 1998); DNA damage induced senescence (DDIS) which can be induced in a number of ways including UV and ionising irradiation or the use of compounds such as etoposide, leading to constitutive activation of the DNA damage response (DDR) and the expression of cell cycle inhibitors; oncogene induced senescence (OIS) occurring through the aberrant activation of oncogenes such as RAS or BRAF; secondary paracrine bystander senescence (BYS) in which neighbouring cells become senescent in response to secreted factors from primary senescent cells; senescence induced through chromatin remodelling (CR); the breakdown of the nuclear barrier leading to nuclear barrier induced senescence (NBIS); Notch induced senescence (NIS) through ectopic NICD activation as well as Ras and Notch (combined) induced senescence (RNIS) (Hoare et al., 2016); OSKM-induced senescence as a by-product of trying to induce pluripotency; induction of senescence through the disruption of ribosomal function (RiboMature) (Pantazi et al., 2019); depletion of deoxyribonucleotide triphosphates (dNTP) induced senescence (Buj et al., 2019); senescence induction through mitotic skipping through the inhibition of CDK1 and MDM2 (MitoSkip) (Johmura et al., 2014); inhibition of the proteasome with drugs such as bortezomib, leading to proteasome inhibition-induced premature senescence (PIIPS) (Skea et al., 2023); highly concentrated treatment with trehalose (Trehalose) (Muto et al., 2023). Control cells could be proliferating or quiescent, and some lines were immortalised or treated with agents that immortalised them as part of the study. Twenty studies compared senescent cells to cells immortalised primarily through hTERT activation, although one study used immortalised cells with p53 knockout (Purcell et al., 2014).

Some comparisons included treatments such as sh/siRNAs against genes designed to observe their effects on senescence, while others used cells from patients with diseases such as breast cancer (Chan et al., 2016), non-small-cell lung cancer (Martinez-Zubiaurre et al., 2013), and Li-Fraumeni syndrome; an inherited syndrome causing vulnerability to rare cancers (Malkin, 1993), here due to mutation of p53 (Purcell et al., 2014). Overexpression of the mitochondrial related gene Parkin (Correia-Melo et al., 2016; Victorelli et al., 2023; Vizioli et al., 2020) also affected gene expression. Another study looked at senescent cells treated with compound ‘1201’, an alcoholic extract from the plant *Solidago alpestris,* that had unknown effects on gene expression (Lämmermann et al., 2018).

To make the database widely accessible, we created a website allowing users to filter for multiple variables to find studies and genes of interest. Users can identify all study or gene data meeting these criteria. For example, comparisons that meet multiple criteria such as ‘OIS in skin fibroblasts with p53 inhibition versus proliferating controls’ can be made using the online database available at: https://research.ncl.ac.uk/cellularsenescence. The median LogFC and p values can also be calculated at the click of a button, and the database can be downloaded for further analysis. The website comes with an “About” page that explains further details.

### Comparison of Senescence Profiles and Biomarker Identification

In our initial analysis we included only the 220 comparisons between senescent cells and proliferating controls without genetic abnormalities or treated with agents that altered gene expression (outside of genes such as RAS, RAF and RCC1 used to induce senescence). Of the 220 comparisons of senescence vs proliferating controls, 196 of them involved REP, DDIS, OIS, or BYS. We therefore compared these four types of senescence to see which genes were commonly significantly different to proliferating cells. For this calculation we used the inverted p value (p_i_ value) (see Methods) so that genes that showed repeated significant change including both increases and decreases compared to proliferating cells, were not counted as genes that showed significant change in a consistent direction. The median value was calculated for each gene for each senescence inducer, and a Venn diagram was plotted showing which genes had significant values for which groups (**Figure 2A**). 28 genes were identified as significant for all four types of senescence; 14 genes consistently downregulated, 10 genes consistently upregulated and 4 genes regulation dependent on senescence type (**Figure 2B**). Gene set enrichment analysis (GSEA) of these 28 genes revealed significant suppression of E2F targets and significant activation of genes upregulated by KRAS signalling (**Figure 2C**). The identification of only two pathways enriched across the four types of senescence mainly reflected that BYS cells had fewer significant changes across all BYS studies (a total of 388 significant genes compared to DDIS, OIS and REP which all identify over 4000 significant genes). There were 916 genes showing consistent and significant changes for OIS, DDIS, and REP. As might be expected, GSEA of the 916 genes common to OIS, DDIS and REP showed significant suppression of the mitotic spindle, G2M checkpoint, E2F targets, and DNA repair (**Figure 2D**), all of which suggest inhibition of the cell cycle and surpassing of the DDR. There was additionally significant inhibition of MTORC1 signalling, spermatogenesis, and MYC targets, while there was significant activation of xenobiotic metabolism, KRAS signalling, p53 pathway and hypoxia pathways. Suppression of MTORC1 signalling is interesting as there is a large focus on rapalogues and how inhibition of mTOR can extend lifespan (Weichhart, 2018). Identifying suppression of MTORC1 as significantly enriched in primary senescent cells provides further evidence of senescence being causal in ageing. However, some studies have found that mTOR signalling is required for senescence (for example, PTEN-loss-induced cellular senescence (Jung et al., 2019)), and that inhibition of mTOR with rapamycin can delay senescence progression (Demidenko & Blagosklonny, 2008; Xu et al., 2014).

**Figure 2.**
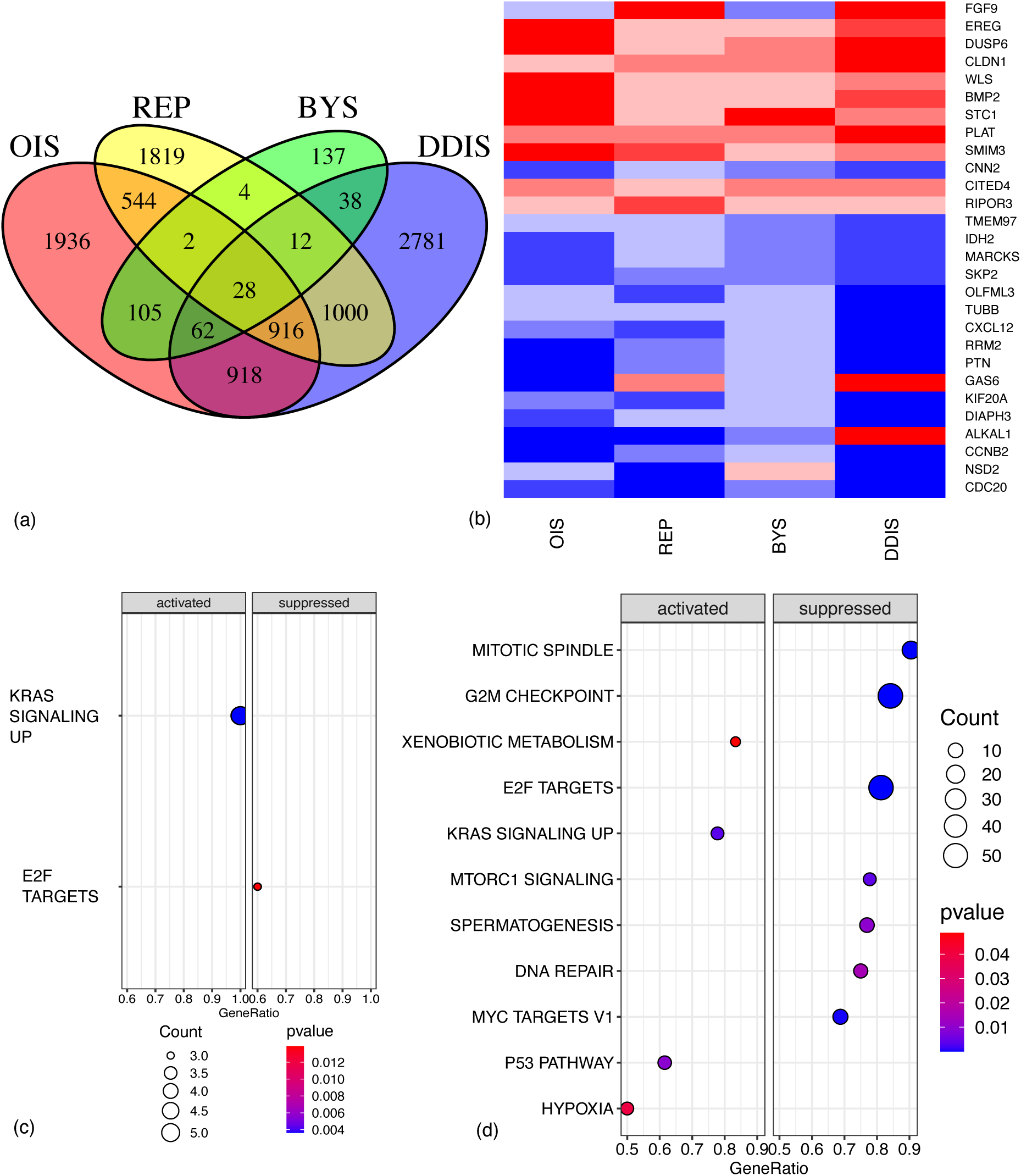
Significant genes and pathways across four types of senescence compared to proliferating control cells. (A) Venn diagram of genes with median p_i_ value that is significant across OIS, REP, BYS and DDIS. (B) Heatmap of the 28 genes that were significant for all four senescence inducers. Upregulated, red; downregulated, blue. (C-D) Dot plot of pathways from GSEA of significantly activated and suppressed pathways identified in (C) the 28 genes common in OIS, REP, BYS and DDIS and (D) 916 genes common in OIS, REP and DDIS. P value refers to the significance of the overrepresentation of the pathway and count reflects the number of genes associated with the pathway. OIS, oncogene induced senescence; REP, replicative senescence; BYS, bystander induced senescence; DDIS, DNA damage induced senescence; GSEA, gene set enrichment analysis.

Similarly, GSEA for the 918 genes showing significance for both OIS and DDIS indicated these pathways plus the activation of TNFA signalling via NFKB and the suppression of genes involved in downregulating the UV response (**Figure S2A**). The analysis strongly suggested that the common changes in expression for senescent cells, with the exception of BYS cells, is the suppression of the cell cycle and DDR, with different inducers suppressing different genes within these pathways. Not all gene set signatures identified significantly enriched pathways; even with a less stringent p value (p <0.01) the 1000 genes significant for both DDIS and REP and the 137 gene significant for just BYS identified no enriched pathways. When disregarding the p value, the 1000 significant genes common to both DDIS and REP identified pathways similar to above such as suppression of E2F targets and the G2M checkpoint (**Figure S2B**), however the 137 genes specific to BYS do not find either of these pathways suppressed when the p value is disregarded (**Figure S2C**). The fact that BYS cells are not suppressing these genes is interesting. Of the four senescence inducers, BYS had the fewest studies and comparisons, which increases the impact of outlier studies when calculating the median p_i_ value. Thus, this difference may simply reflect that BYS cells have less data available. However, it may also reflect the difference between primary and secondary senescence which is still being understood (Admasu et al., 2021; Teo et al., 2019).

Notably, in their non-systematic review of transcriptomic data, Hernandez-Segura et al. (2017) identified a 55 gene core signature for all types of senescence observed. The analysis included six different fibroblast strains (BJ, IMR90, HFF, MRC5, WI38, and HCA-2) for three different inducers (REP, OIS, and DDIS). None of our 28 genes with significant median p_i_ values were in this 55 gene core signature, and only CNTLN, FAM214B, MEIS1, PLK3 and TSPAN13 were consistent with our 916 genes which excluded BYS. Another study by Casella et al. (2019) produced a 68 gene core signature for cellular senescence (not including BYS), of which only one gene (CLDN1) was consistent with our 28 core gene signature, which was upregulated in both cases. There were sixteen genes consistent with our 916 gene signature excluding BYS: ANP32B, CDCA7L, DHRS7, ELMOD1, HIST1H1A, HIST1H1D, HIST2H2AB, ITPRIPL1, JCAD, KIAA1671, LBR, LRP10, PAM, PARP1, PTMA, and SLC9A7. Only POFUT2 was consistent between the core signatures identified between Casella et al. (2019) and Hernandez-Segura et al. (2017). Notably both studies included non-fibroblast cells, and Hernandez-Segura et al. (2017) only included genes that were also significantly different to quiescent cells. However, the identification of a consistent transcriptional biomarker for senescence is clearly problematic. One possibility for the inability to identify any clear core senescence signature is that the senescence profile may change temporally and our above data spans from day 0 to day 112 post-senescence induction, while Hernandez-Segura et al. (2017) and Casella et al. (2019) only investigate up to day 10 post-senescence induction. To support this systematic analysis, we investigated temporal changes in gene expression of core senescence proteins and developed a qualitative computational model of DDIS and OIS which simulates temporal changes of protein expression as senescence progresses, demonstrating how different stages of senescence are characterised by different signatures.

### Cellular senescence protein network and phenotypes

The senescence model network was developed based on a non-systematic literature review of protein profiles and our systematic transcriptomic analysis to determine the temporal senescence phenotype. From these combined reviews we developed a total of 12 molecular profiles defining senescence or specific to the subtypes, four of which were related to how senescent cells responded when key network components were knocked down (**Table 3**). A multiple sub-model approach was taken to improve the understanding of connections between proteins, ensuring they were biologically correct and not just modelling artefacts.

**Table 3.** Phenotypes of cellular senescence. *0 denotes no studies can be found related to this phenotype at the protein level; 1 denotes limited published studies support this described phenotype at the protein level; 2 denotes multiple studies support this described phenotype at the protein level; 3 denotes this is a well-established protein phenotype in senescent cells. † 4 days post-senescence induction correlated with arbitrary time 60 in model simulations. †† 5-8 days post-senescence induction correlates with arbitrary time 65-85 in model simulations. Y, yes; N, no

| Senescent cell phenotype | How established is this phenotype at the protein level * | Is this phenotype reflected in the transcriptomic systematic analysis? | Do model simulations match the senescent cell phenotype criteria? |  |  |
| --- | --- | --- | --- | --- | --- |
|  |  |  | Sub-model A | Sub-model B | Model C |
| p21 and p16 are still expressed once senescence has been induced for more than 4 days† | 3 | Y | Y | Y | Y |
| There is higher p53 activation and p21 expression in DDIS than in OIS | 0 | Y | N | N | Y |
| There is higher p16 expression in OIS than in DDIS | 1 | Y | Y | Y | Y |
| Expression of p16 increases later in DDIS than OIS | 0 | Y | Y | Y | Y |
| Changes in Notch signalling activity is temporally associated with a switch in the SASP profile | 1 | Y | N | N | Y |
| There is a stronger inflammatory phenotype in OIS than in DDIS | 2 | Y | N | Y | Y |
| The inflammatory phenotype begins to be established on days 5-8 after senescence induction†† | 3 | Y | N | N | Y |
| There is more p38 phosphorylation in OIS than in DDIS | 1 | N | Y | Y | Y |
| **KNOCKDOWN CRITERIA** |  |  |  |  |  |
| Knockdown of p53 results in upregulation of p16 expression | 0 | Y | N | N | Y |
| Knockdown of p53 results in decreased p21 expression | 3 | Y | Y | Y | Y |
| Knockdown of p53 results in upregulation of p38 in DDIS | 3 | Y | N | N | Y |
| Knockdown of RELA results in decreased p53 signalling | 0 | Y | N | Y | Y |

The first model developed, sub-model A (**Figure 3A**), is composed of cell cycle arrest and DDR proteins such as p53 and p21, in addition to RAS and p38. These were included as a source of senescence stimulus. Both DDIS and OIS are known to need overwhelming activation of the DDR leading to cell cycle arrest (Kumari & Jat, 2021), with hyperactivation of an oncogene such as RAS additionally required for OIS (Liu et al., 2018). This provided a foundation for constructing a network which would be representative of cellular senescence. To build upon sub-model A, sub-model B (**Figure 3B**) was developed with the addition of inflammatory SASP proteins. The SASP is a key characteristic of senescent cells therefore the ability for a model to recapitulate SASP dynamics is essential. As observed in the literature the SASP begins to be established 5-8 days after senescence induction (Coppé et al., 2010). Lastly, as the SASP is known to not be immediately expressed upon senescence induction a method to modulate SASP induction was required in the model. In model C (The Senescence Induction Model, **Figure 3C**), Notch signalling was introduced. Dynamic changes in Notch signalling activity have been shown to modulate SASP activity (Hoare et al., 2016), therefore by introducing an event termed ‘The Notch Switch’ we were able to dynamically change Notch signalling at specific timepoint, allowing for modulation of the SASP.

**Figure 3.**
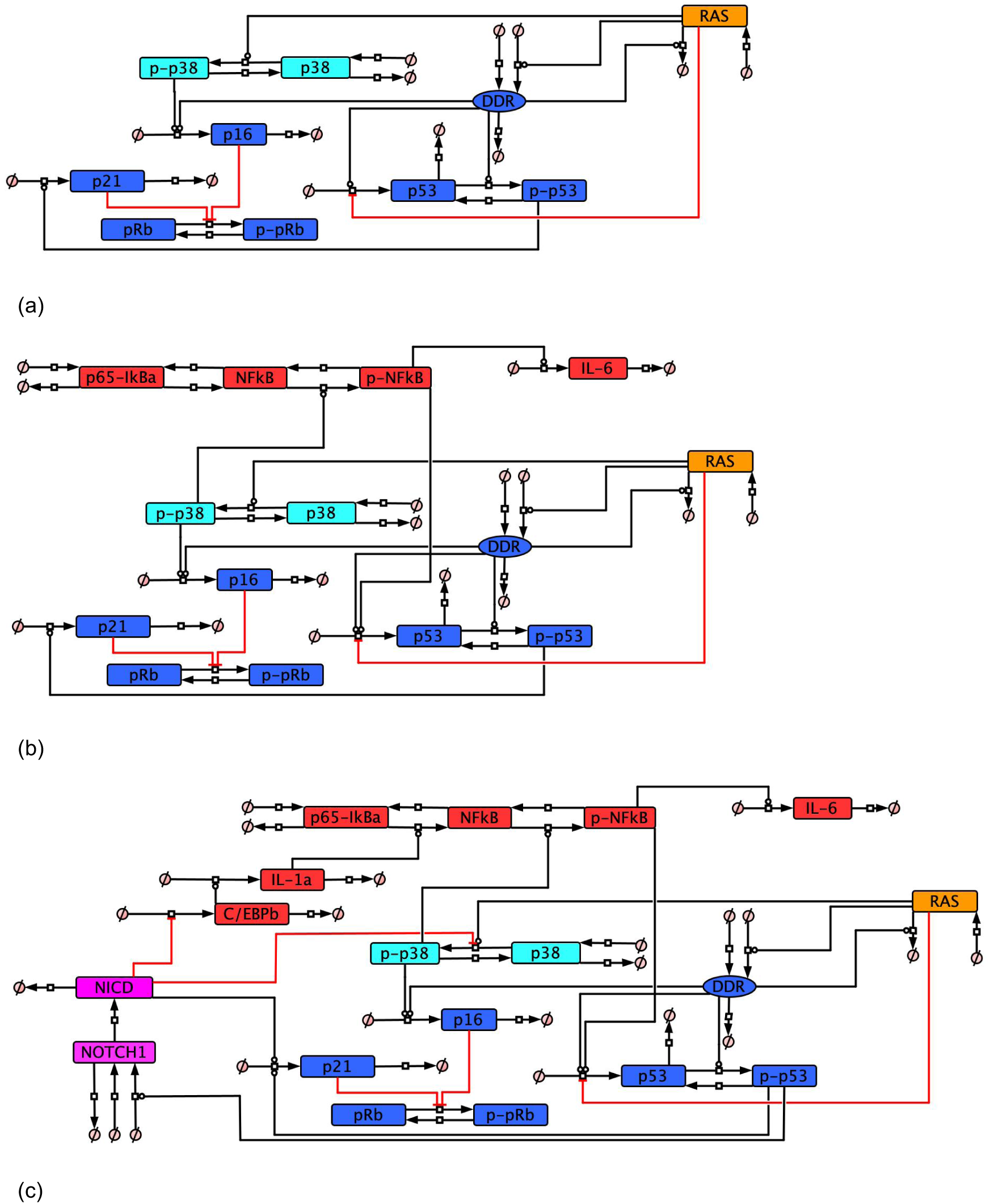
Cellular senescence protein network. (A-C) Development of protein networks to meet the devised cellular senescence phenotype criteria points. All interactions in the network have been non-systematically searched in the literature, providing evidence to justify the network (**Table S2**).(A) sub-model A; (B) sub-model B; (C) model C – the Senescence Induction Model.

In all models, DDIS was simulated by introducing an input which would simulate a large amount of DNA damage through a high activation of the DDR, enough to induce cell cycle inhibitor expression and therefore arrest. We simulated OIS by inducing high level RAS activity which led to activation of p38/p16 and subsequently DDR activation and cell cycle arrest. The model can be adapted to investigate different dynamic aspects of senescence such as the impact of changes in Notch signalling (Hoare et al., 2016; Teo et al., 2019) or the impact of p53 KD or inhibition (Kumari et al., 2021; Zhang et al., 2021; Zhu et al., 2022).

### Activity of p53 in senescent cells

Our study and those by Hernandez-Segura et al. (2017) and Casella et al. (2019) indicate that the standard markers of senescence, including those believed to be causal in senescence induction such as p53 and p21, are not reliable biomarkers. Therefore, we looked more deeply at the genes commonly associated with senescence, attempting to identify conditions where they were demonstrably and reliably active or upregulated at the mRNA level.

We first looked at DDR genes thought to play a central role in initiating the senescence response. Double strand breaks or uncapped telomeres activate ATM and ATR followed by downstream CHEK1 and CHEK2 which activate p53 and cause the transcription of p21. Splitting the data up by timepoint into groups (0-4 days, 5-7 days, 8-11 days, 12-14 days, and 15+ days), we looked at these damage response genes. Evidence for 12 to 14 days, excluding REP, is limited to eight comparisons from seven studies, while the 15+ day data is limited to five comparisons from four separate studies only investigating DDIS. Further research is required at these late time points if consensus is to be reached on the whole transcriptomic profile. The 15+ day data is not plotted due to limited datapoints. REP cells were also included split into two categories: 0-40 day post-senescence induction and 41+ days post senescence induction. As the vast majority of studies of REP cells did not state the timepoint after induction, these were put in the 0-40 day group under the assumption that waiting 41+ days reflected a deliberate attempt to look at the longterm senescence gene profile.

ATM (**Figure 4A**) and ATR (**Figure S3A**) mRNAs showed no observable trend in LogFC, which stayed around the level of proliferating cells, although ATM expression in DDIS was higher than proliferating cells between days 12-14 post-senescent induction. The same was true for CHEK1 (**Figure 4B**) and CHEK2 (**Figure S3B**), except that CHEK1 was observably reduced compared to proliferating cells in both DDIS and OIS at least until day 12. CHEK1 and CHEK2 activity is primarily increased by phosphorylation by ATM and ATR (Ahn et al., 2000; Jazayeri et al., 2006). A function of both CHEK1 and CHEK2 is to phosphorylate CDC25A, causing its degradation. The mRNA data suggest that CDC25A was decreased compared to proliferating cells (**Figure S3C**), potentially sufficient to induce the S and G2 checkpoints (Falck et al., 2001; Xiao et al., 2003). Additionally, CDC25A expression was significantly lower in DDIS compared to OIS at days 0-4 (p value < 0.01) and days 8-11 (p value < 0.05). However, the main role of CHEK1/2 is thought to be in the stabilisation of p53 (Chehab et al., 2000). Notably, this is not reflected in the transcriptional profile of p53, which shows no evidence of an increase in mRNA compared to proliferating cells, and possibly a decrease at some time points (**Figure 4C**). Not observing a change in p53 mRNA expression likely reflects the pulsatile signalling of p53 (Hunziker et al., 2010; Sun et al., 2011), which is bound and inactivated by MDM2 targeting it for ubiquitin-mediated degradation (Michael & Oren, 2003). Although p53 activity is regulated in large by post-translational modifications and coactivators (Fielder et al., 2017), it is also a short-lived protein, and must in some way be regulated at the transcriptional level; however, the pulses are likely too fast for a single measurement or measurements across multiple days to capture the average level of p53 mRNA compared to control cells (Hunziker et al., 2010; Sun et al., 2011).

**Figure 4.**
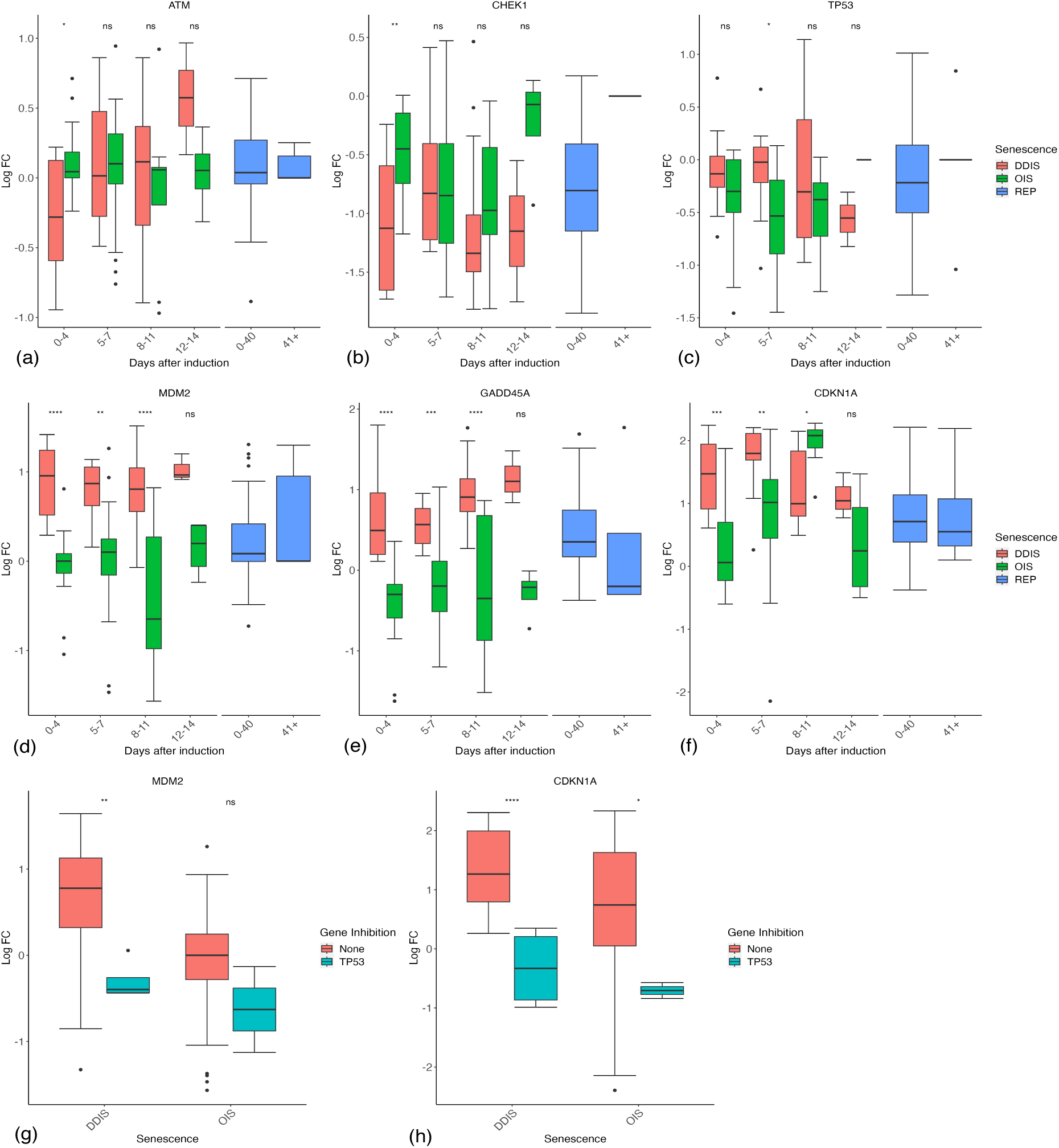
Expression of the damage and p53 response in senescent cells. (A-F) Gene expression during the timeline of senescence induction measured in days after the initial stimulus. (G-H) Gene expression for different senescence inducers with and without p53 inhibition. Control groups for inhibition include all data for days 1-11. DDIS, DNA damage induced senescence; OIS, oncogene induced senescence; REP, replicative senescence; LogFC, log fold change; p value refers to significance in expression between DDIS and OIS (A-F) and expression with and without gene inhibition (G-H), * p value < 0.5; ** p value < 0.01; *** p value < 0.001; **** p value < 0.0001.

Notably, the level of mouse double minute 2 (MDM2) mRNA, the negative regulator of p53, shows an observable increase in DDIS (**Figure 4D**). Interestingly, expression of MDM2 is significantly higher in DDIS cells compared to OIS cells at 0-4 days (p value < 0.0001), 5-7 days (p value < 0.01) and 8-11 days (p value < 0.0001) post-senescence induction. Although MDM2 inhibits p53, increased expression reflects increased p53 activity, as p53 induces the transcription of MDM2 (Barak et al., 1993). This trend is reinforced by other p53-induced genes such as GADD45A (Kastan et al., 1992) and p21 (*CDKN1A*) (**Figure 4E-F**), which are increased across all timepoints in DDIS, while only p21 is notably increased in OIS. Notably, the systematic analysis gives little indication that p53 activity decreases before day 11, with p21, GADD45A, and MDM2 trending to increase at 8-11 days compared with 5-7 days in both OIS and DDIS, and typically be above proliferating controls at days 12-14 days post-senescence induction.

To confirm the role of p53 in the upregulation of these genes, we looked at the studies which inhibited p53. REP cells were excluded as these cells have no defined time after senescence induction, as was one comparison at day 28, long after p53 signalling is thought to have subsided (Robles & Adami, 1998). As expected, p53 mRNA was observably reduced in p53 inhibition studies (**Figure S4A**). As predicted, the downstream targets of p53, MDM2 and p21 mRNAs, both showed observable reductions in the p53 inhibition group (**Figure 4G-H**), but interestingly this was not true of GADD45A or B (**Figure S4B-C**). We concluded that although p53 mRNA (Hernandez-Segura et al., 2017) was not a reliable biomarker of senescent cells, the combined transcriptional data from all available studies suggest that p53 is highly active in senescent cells up to at least 14 days (**Figure 4D-F**). However, another clear observation is that its activity is lower (as measured by p21, MDM2, and GADD45A) in OIS compared to DDIS, suggesting the p53 may play a larger role in DDIS than in OIS.

p53 signalling is known to be active in senescence (Mijit et al., 2020; Rufini et al., 2013), and our systematic analysis supports long-lasting p53 signalling in senescence, with stronger p53 signalling in DDIS compared to OIS (**Figure 4D-F**). Furthermore, as seen in published work (Chien et al., 2011), KD of p53 results in diminished p21 expression. In modelling temporal changes of p53, pp53 and p21 at the protein level, simulations of sub-model A do not meet all criteria pertaining to p53 signalling in senescence (**Table 3**) as p53 signalling is lower in DDIS than in OIS (**Figure 5A-C**). This reverse in expected signalling activity is not due to errors in modelling but due to a lack of input from additional proteins which provide p53 with further context on how it behaves in a cell. The introduction of a p53 KD, however, was successful at reducing p53 signalling as expected (**Figure 5A-C**). Sub-model B (which contains the network from sub-model A with the addition of some inflammatory proteins) also does not meet all criteria regarding p53 signalling as although p53 signalling is initially higher in DDIS simulations, expression of these proteins in OIS overtakes expression in DDIS as the simulation progresses (**Figure 5D-F**) which is not seen in the literature or in our transcriptomic analyses (**Figure 4D-F**). This suggested that there were further proteins required to add biological context to simulations. Simulations of the Senescence Induction Model meet all criteria related to p53 signalling as there is sustained higher activation of p53 signalling in DDIS compared to OIS (**Figure 5G-H**); p21 is expressed in late senescence (**Figure 5I**); KD simulations are able to recapitulate reduced p53 signalling when a p53 KD was introduced (**Figure 5I**); reduced p53 signalling is observed in OIS and DDIS when a RelA KD is introduced (**Figure 5G-I**). As the Senescence Induction Model can recapitulate all criteria related to p53 and p21, this suggests the network is complex enough so that true biological interactions can be portrayed but not too simple so that the details are lost.

**Figure 5.**
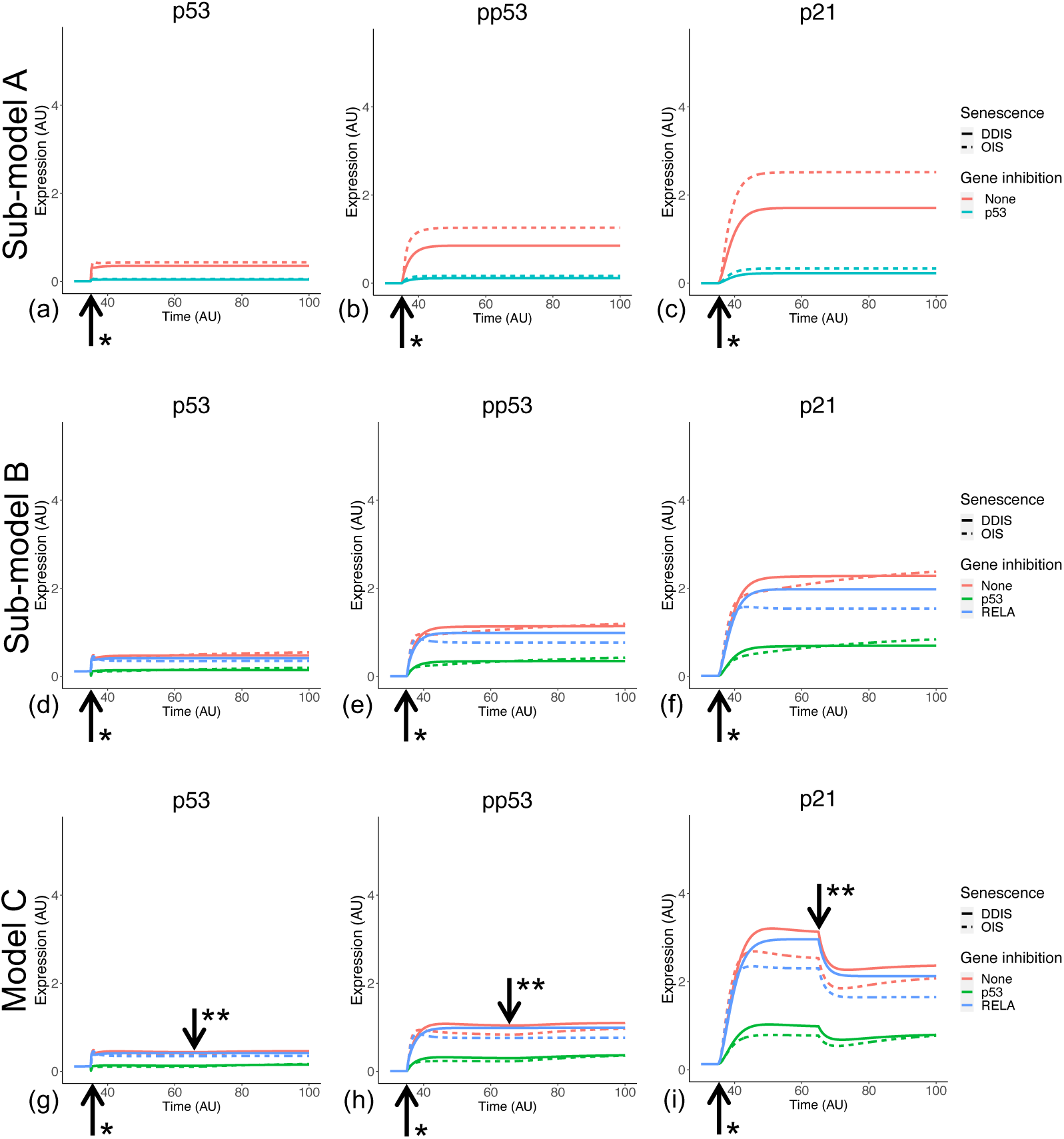
Simulation of p53 signalling dynamics. Simulations show temporal expression of p53 and p21, including the phosphorylation status of p53, in DDIS and OIS and also when a p53 or RELA KD is introduced. Units are arbitrary. *Senescence and/or KD induced. **Notch Switch induced.

### OIS and DDIS rely on different mechanisms for arrest

The differences between OIS and DDIS are still being elucidated. DDIS reflects the direct sub-apoptotic chronic induction of the DDR, typically mediated by double strand breaks (DSBs), but OIS need not. Some studies have shown that OIS is bypassed in the absence of the DDR (Bartkova et al., 2006; Di Micco et al., 2006; Mallette et al., 2007), and RAS-induced OIS cells can re-enter the cell cycle if the DDR is inactivated, reflecting that OIS relies on the DSBs induced by the aberrant activation of oncogenes and the resultant hyperproliferation (Di Micco et al., 2006). However, other reports suggest that OIS can be induced independently of the DDR (Alimonti et al., 2010), although still requiring p53 (Wolyniec et al., 2009) or p16 (Bracken et al., 2007). Interestingly, while p21 is significantly higher in DDIS compared to OIS up to day 11 post-senescence induction (**Figure 4F**), p16 (*CDKN2A*) is significantly higher in OIS compared to DDIS up to day 11 post-senescence induction, and non-significantly higher at days 12-14 (**Figure 6A**). Upon inhibition of p53 in senescent cells, p16 expression does not significantly change (**Figure 6B**), suggesting that p16 is independent of p53 signalling (Alcorta et al., 1996). Although it does look to increase in DDIS but not OIS, suggesting p16 may be inhibited or negatively regulated by p53 signalling.

**Figure 6.**
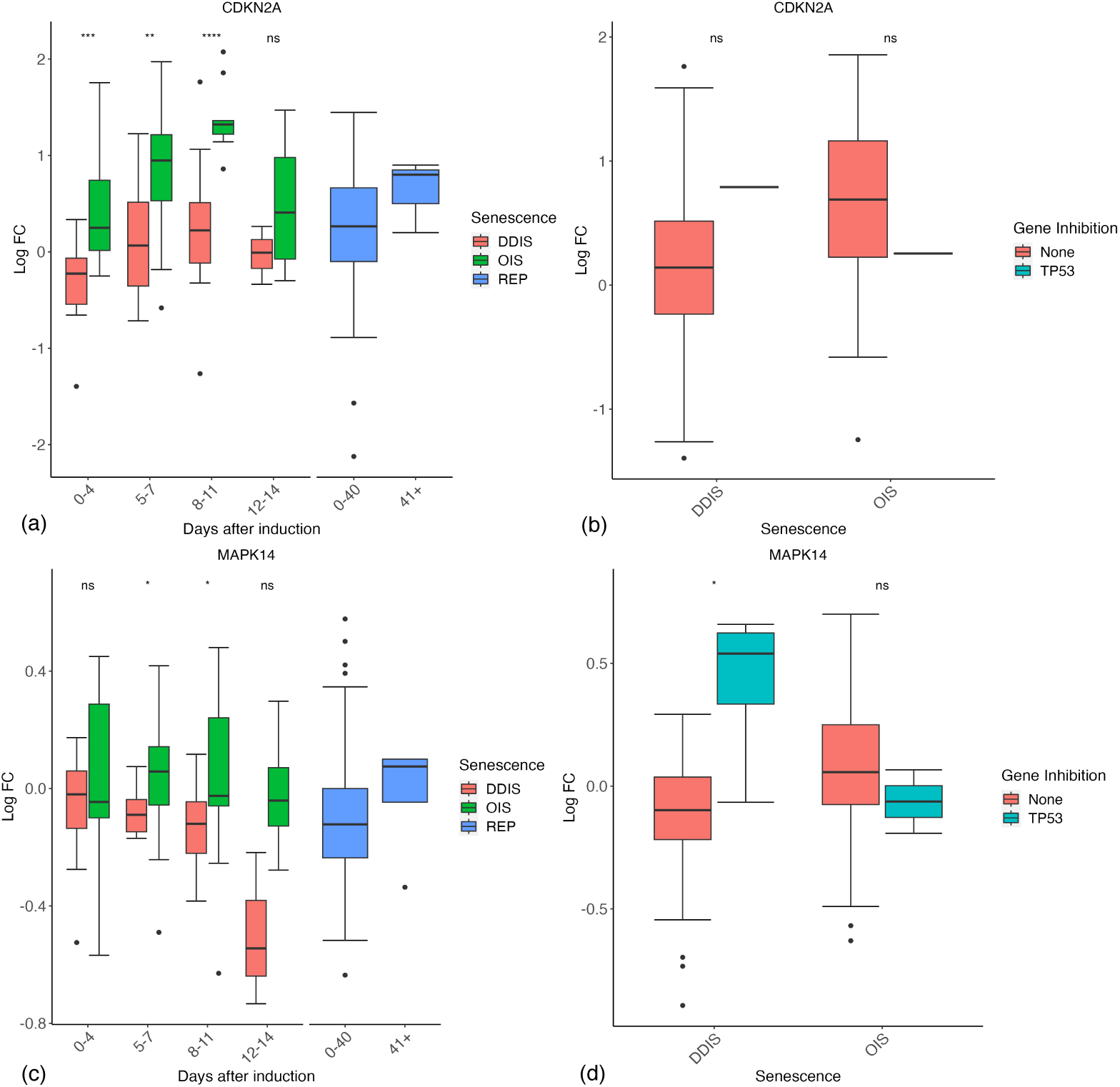
Expression of p16 and p38 genes in senescent cells. (A-B) Expression of p16 (CDKN2A) during different timepoints (A) and with p53 inhibition (B). (C-D) Expression of p38 (MAPK38) during different timepoints (C) and with p53 inhibition (D). Control groups for inhibition include all data for days 1-11. DDIS, DNA damage induced senescence; OIS, oncogene induced senescence; REP, replicative senescence; LogFC, log fold change; p value refers to significance in expression between DDIS and OIS (A & C) and expression with and without gene inhibition (B & D), * p value < 0.5; ** p value < 0.01; *** p value < 0.001; **** p value < 0.0001.

The activation pathway of p16 is still somewhat controversial. One suggestion is that DNA damage activates p38 (Bulavin et al., 2003; Ito et al., 2006; Iwasa et al., 2003), which then activates p16 (Spallarossa et al., 2010). Our data are consistent with this idea, with p38 (*MAPK14*) being higher in OIS than DDIS, significantly higher 5-11 days post-senescence induction (p value < 0.05) (**Figure 6C**), and higher in p53 inhibited DDIS cells (**Figure 6D**). Consistently, Freund et al. (2011) found that p53 inhibited p38 phosphorylation which has implications for the SASP. Several genes showed a stronger response to p53 inhibition in DDIS than in OIS, presumably reflecting that p53 activity is higher in DDIS.

It has been repeatedly suggested that p21 is transient in senescence, required only for induction (He & Sharpless, 2017; Kumari & Jat, 2021). Robles and Adami (1998) showed p21 levels were decreasing by day 8 in DDIS, suggesting that p53 activity peaked around day 4. While this may be true, we saw little evidence of mRNA decline by 8-11 days (**Figure 4F**), and even at days 12-14 certainly in DDIS expression of p21 mRNA was still higher than in control proliferating cells. Two other frequently cited studies describing transient p21 levels are in REP, and show p21 levels declining over weeks of passaging (Alcorta et al., 1996; Stein et al., 1999). Our systematic analysis indicates p21 levels are still increasing at 8-11 days compared to 5-7 day cells. From this data, p21 seems no more transient than p16, which is generally described to be absent in early senescence and rise slowly over time.

Robles and Adami (1998) showed p16 mRNA was no higher than control at day 4 DDIS, slightly increased at day 12 and then peaking at day 30. Stein et al. (1999) indicated that in REP, p16 began its steepest increase after 30 weeks of passaging (compared to 10 weeks for p21). However, systematic analysis indicates that p16 is already increasing in OIS by day 4, and is significantly higher in OIS than DDIS (p value < 0.001) (**Figure 6A**). The rise is less obvious in DDIS, but it does tend to follow an increase temporally.

This contrasts p21, which rises at 0-4 days in DDIS but median expression does not increase above 0 LogFC until 5-7 days in OIS. Speculatively, this may reflect that the damage is the primary initiator in DDIS, promptly activating p21, whereas in OIS the damage from hyperproliferation may take longer, while RAS, p38, or other mechanisms independently activate p16. In DDIS, the rise in p16 is slower as reflected by the median LogFC just above zero even by 8-11 days. However, (Hoare et al., 2016) show western blots of p16 protein levels increasing by 2 days for both OIS and DDIS. The band at 8 days is observably thicker for OIS than in DDIS, which is again consistent with the systematic analysis. Notably, while p38 may activate p16, p38 phosphorylation is increased between 6-8 days in OIS (Freund et al., 2011), so it is unlikely to explain the early rise, but if p38 is also activated by RAS signalling (Chen et al., 2000), it may reflect an additional mechanism upregulating p16 in OIS but not DDIS, which might explain the later difference.

Although there is controversy surrounding p16 dynamics in senescence, our analyses suggest p16 expression is maintained into late senescence, significantly more expressed in OIS compared to DDIS, and is expressed in OIS before it is expressed in DDIS (**Figure 6A**). Our systematic analysis reveals limited change in p38 expression as senescence progresses (**Figure 6C**), something which is recapitulated at the protein level in the Freund et al. (2011) study. Freund et al. (2011)(Freund et al., 2011) also demonstrate that while total p38 levels remain similar throughout senescence, phosphorylation levels increase from day 4 to day 6 and continue to increase up to day 10 which was the final timepoint analysed. As phosphorylation status cannot be explicitly determined at the mRNA level, we are unable to check if this is reflected in the transcriptomic data. However, phosphorylation can be investigated in the computational model, further demonstrating the need for an integrated approach to investigating complex mechanisms such as cellular senescence.

Sub-model A simulates all criteria pertaining to the normal senescence phenotypes related to p16 and p38 (**Table 3**) as p16 expression is maintained throughout senescence and expression of p16 begins earlier and is more highly expressed in OIS than in DDIS (**Figure 7A**). Furthermore, there is more phosphorylated p38 (pp38) in OIS than in DDIS simulations (**Figure 7B-C**). However, when p53 is knocked down, there is only one coloured line for DDIS and OIS in simulation plots, indicting no change in p16, p38 and pp38 expression (**Figure 7A-C**). Likewise, sub-model B meets all normal senescence phenotype criteria but does not meet any of the KD criteria (**Figure 7D-F**), demonstrating that proteins not included in the network of sub-model B are required for correct functioning of p16 and p38 in senescence. The Senescence Induction Model meets all normal senescence phenotypes and KD phenotypes relating to p16 and p38 as expression of p16 is higher and begins earlier in OIS than in DDIS and is increased when p53 is knocked down (**Figure 7G**) as expected from our analyses (**Figure 6A**). One study does find decreased p16 transcript expression when p53 is knocked down (Georgilis et al., 2018), however this could be due to study design and may not reflect protein level changes. p38 expression is steady throughout simulations of DDIS and OIS (concurrent with observations in our analyses (**Figure 6C**) and published protein level studies (Freund et al., 2011)) while pp38 increases, particularly in late senescence, and KD of p53 results in increased p38 expression (**Figure 7H-I**), which is seen at the transcript (**Figure 6B & D**).

**Figure 7.**
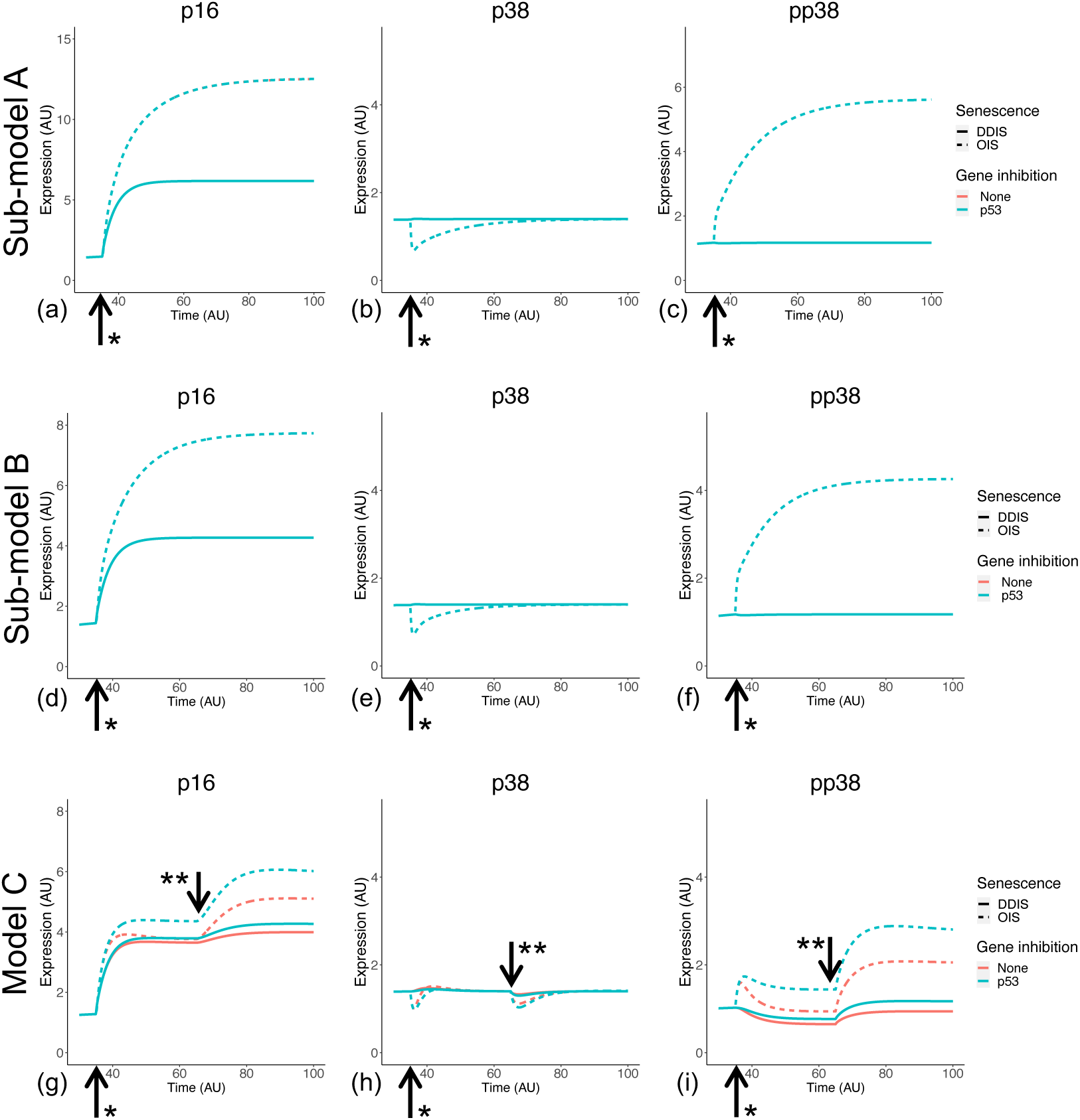
Simulation of p16 and p38 in senescence. Simulations show temporal expression of p16 and p38 proteins, including phosphorylation status of p38, in DDIS and OIS and when a p53 KD is introduced. Units are arbitrary. *Senescence and/or KD induced. **Notch Switch induced.

### Induction of the SASPs

The processes that lead to the induction of the SASP are still uncertain. As discussed by Freund et al. (2011), one possible mechanism is that p38 activates NF-κB. Perhaps the most detailed temporal profile comes from Hoare et al. (2016) who suggested that the initial SASP of OIS and DDIS was a TGFβ-rich secretome, which due to a breakdown in Notch signalling around day 4-5 became an inflammatory secretome.

Notch signalling is a juxtacrine mechanism. When a ligand from one cell binds the Notch receptor of an adjacent cell, there are multiple cleavage events which results in the release of the Notch intracellular domain (NICD) which can then translocate into the nucleus and enact Notch targeted gene transcription (Bray, 2006). As Notch signalling is regulated through receptor cleavage and ligand binding, we would not expect to see much change in expression of the Notch1 receptor gene. Analysis of Notch1 in our database did not reveal much other than that expression is higher in OIS than in DDIS, significantly higher at day 0-4 (p values < 0.01) and 8-11 days (p vales <0.001) (**Figure 8A**). We then used the database to investigate changes in Notch signalling through expression of two well-known Notch target genes, HES1 and HEY1. It should be noted that Notch1 transcriptional activity is still not well understood, even HES1 is not always responsive to Notch1 activation (Kopan & Ilagan, 2009; Lee et al., 2007). In our analyses, both HES1 and HEY1 show an increase in median expression compared to control proliferating cells at 0-4 days in DDIS and OIS. Expression stays relatively similar at days 5-7 for HEY1 (**Figure 8B**) before decreasing at days 8-11, while median HES1 expression decreases from days 0-4 to days 5-7 (**Figure 8C**). Notably, this corresponds with the study by (Hoare et al., 2016) at the protein level who see an initial increase in Notch signalling activity (determinised via the presence of the Notch intracellular domain (NICD) and Hes1) followed by a decrease between days 4-6.

**Figure 8.**
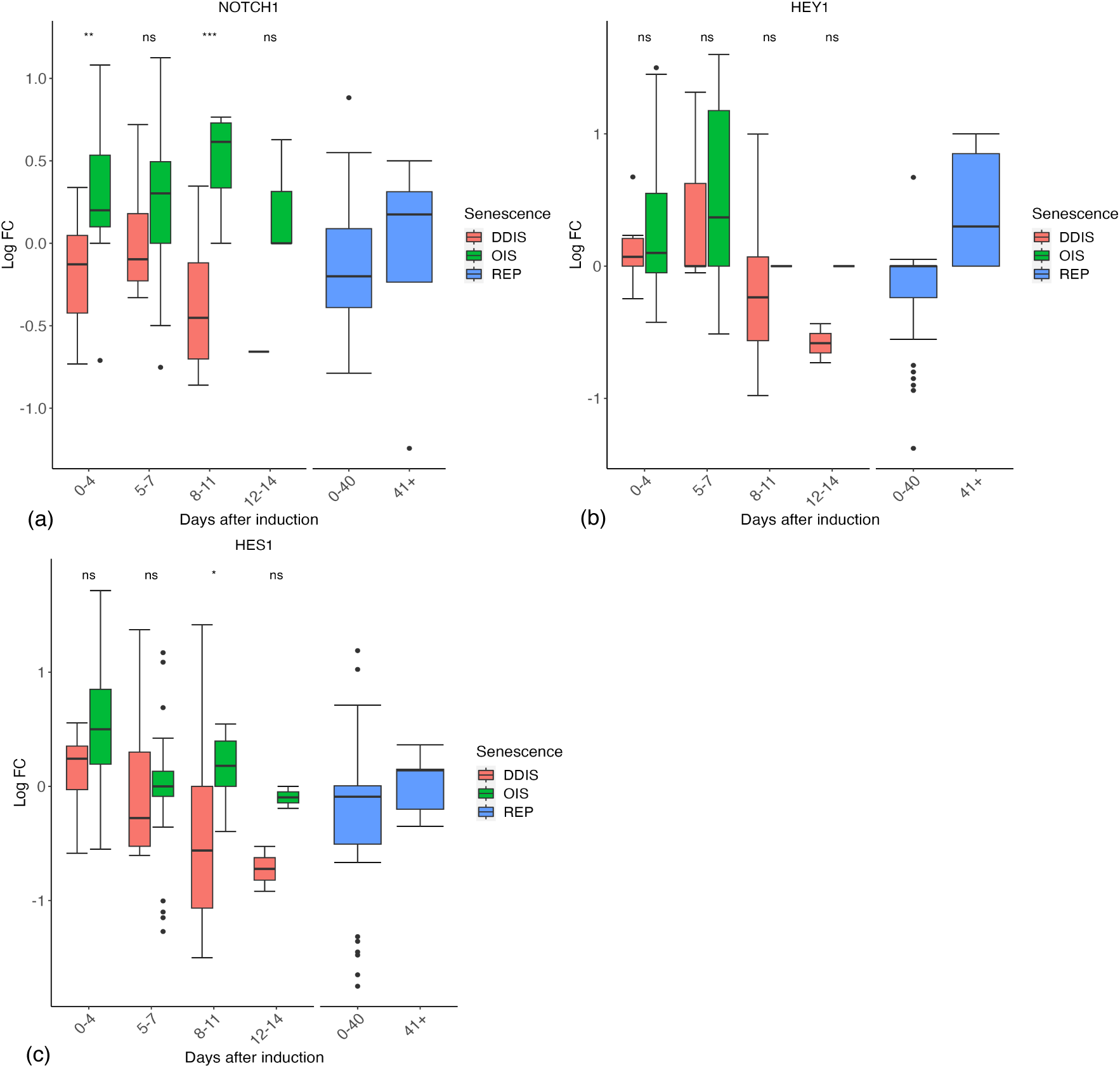
Expression of Notch signalling related genes in senescent cells. (A-C) Gene expression during the timeline of senescence induction measure in days after the initial stimulus. DDIS, DNA damage induced senescence; OIS, oncogene induced senescence; REP, replicative senescence; LogFC, log fold change; p value refers to significance in expression between DDIS and OIS,* p value < 0.5; ** p value < 0.01; *** p value < 0.001; **** p value < 0.0001.

To model a change in Notch signalling dynamics, we introduced the NOTCH1 receptor and NICD as protein species into the Senescence Induction Model. Neither sub-model A or B have Notch proteins present. This model is at the single cell level therefore to induce a change in Notch receptor activity we introduced an event termed the Notch switch which would result in the downregulation of NICD expression, representing less ligand-receptor activity as less NICD is cleaved from NOTCH1.

Simulations of the Senescence Induction Model demonstrate dynamic changes in Notch signalling temporally. NICD is expressed in early senescence while senescence is being established, indicating active Notch signalling. This is followed by a reduction in NICD and therefore Notch signalling once the Notch switch is activated (**Figure 9**). Simulation dynamics recapitulate published protein profiles in senescence (Hoare et al., 2016) and reflect our systematic analysis of HES1 transcript expression (**Figure 8C**), supporting the observation that dynamic Notch signalling is involved in regulation of the SASP.

**Figure 9.**
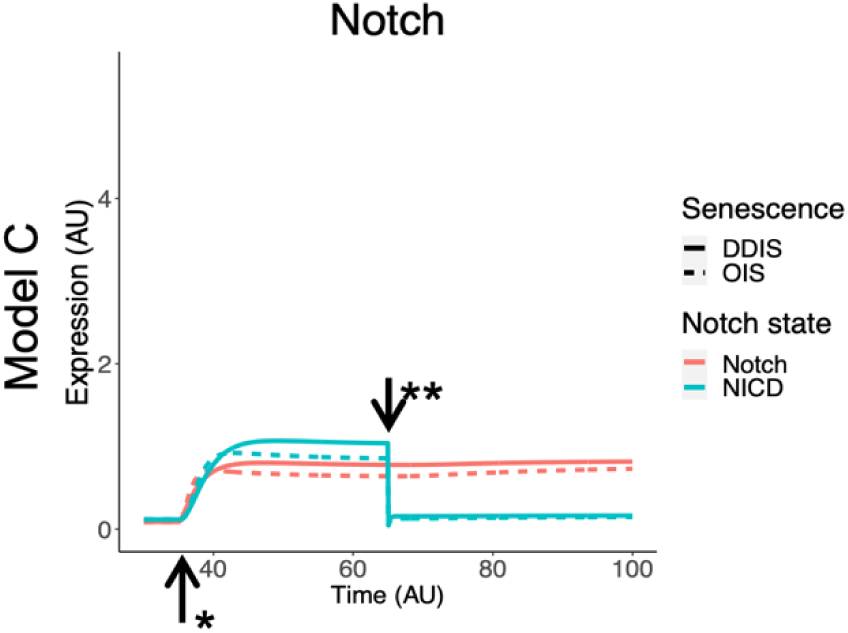
Simulation of Notch signalling dynamics in cellular senescence. Simulations show dynamic temporal changes in Notch signalling from active to inactive in both DDIS and OIS. Units are all arbitrary. *Senescence induced. **Notch Switch induced.

### The SASPs of OIS and DDIS are governed by the activity of p38 and p53

NF-κB is essential for the production of the inflammatory SASP (Chien et al., 2011; Freund et al., 2011), and co-suppression with p53 leads to bypass of arrest. Notably, BJ fibroblasts required only shRNA against p65 to bypass arrest, which the authors concluded may reflect the previously identified less robust senescence program in this cell type (Beauséjour et al., 2003). We compared gene expression for the different cell lines, discussed in **Figure S5**. The results were consistent with a different profile for BJ cells, however this was not investigated in depth here.

Importantly, there is strong evidence of an increased inflammatory response in OIS compared with DDIS, with the main SASP factors including IL6, IL8 (*CXCL8*), and IL1B all showing higher levels in OIS cells over DDIS at least between 5-11 days (**Figure 10A-C**). Although this is not significant at any timepoints, the median expression is typically higher in OIS than in DDIS. Interestingly, in REP cells the expression of IL6, IL8 and IL1B are all increased at days 40+ compared to 0-40 days whose median expression is around 0 LogFC and therefore similar to proliferating control cells (**Figure 10A-C**). This shows that while the SASP may be more strongly expressed in OIS, the SASP is still an important mechanism in other types of senescence.

**Figure 10.**
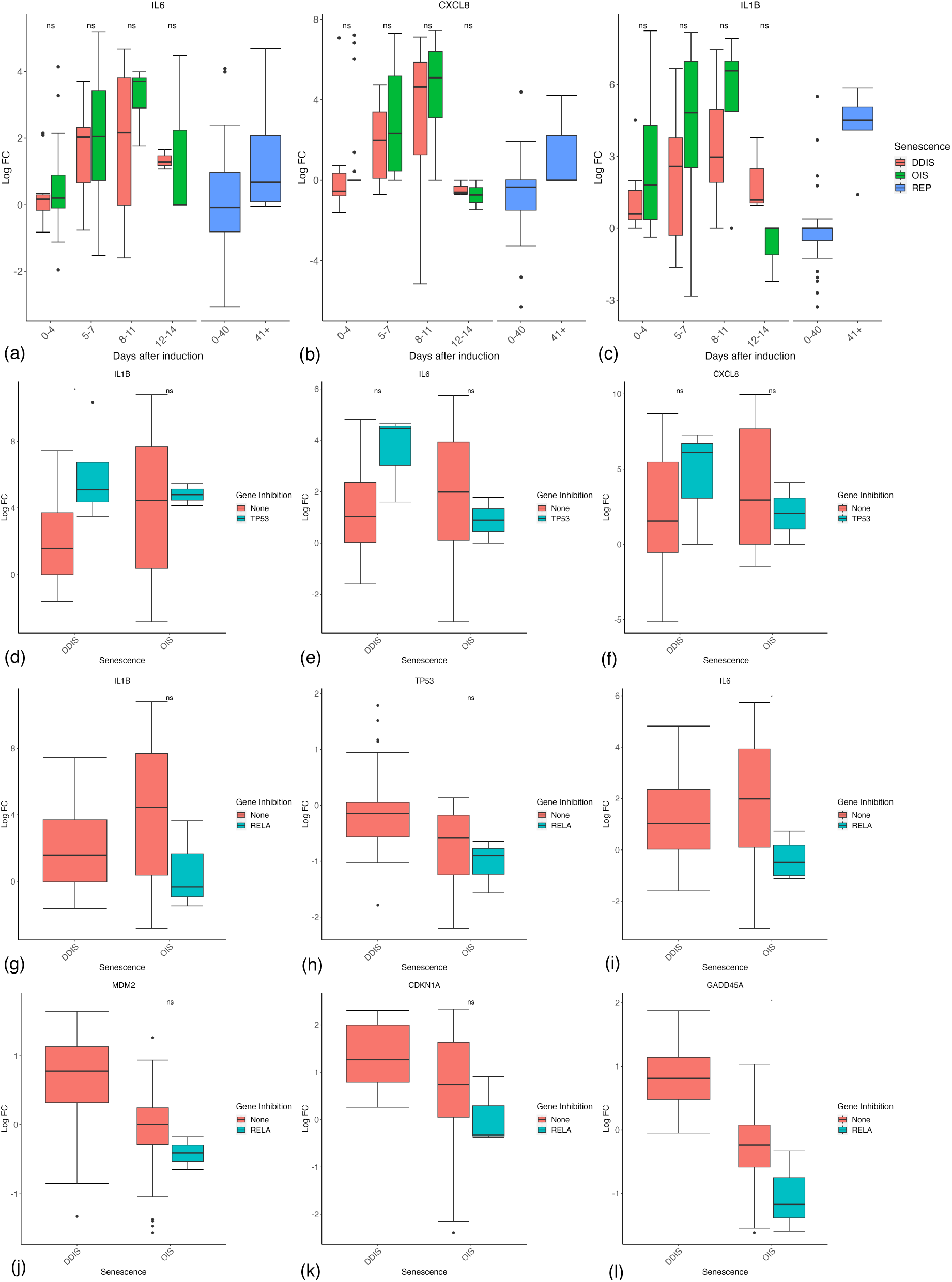
Expression of inflammatory and p53 genes in senescent cells. (A-C) Gene expression during the timeline of senescence induction measured in days after the initial stimulus. (D-E) Gene expression for different senescence inducers with and without p53 inhibition. (F-K) Gene expression for different senescence inducers with and without RELA inhibition. Control groups for inhibition include all data for days 1-11. DDIS, DNA damage induced senescence; OIS, oncogene induced senescence; REP, replicative senescence; LogFC, log fold change; p value refers to significance in expression between DDIS and OIS (A-C) and expression with and without gene inhibition (D-L), * p value < 0.5; ** p value < 0.01; *** p value < 0.001; **** p value < 0.0001.

Increased expression of inflammatory proteins increasing from 5-7 days is consistent with the timing of the SASP concluded by others (Coppé et al., 2008; Freund et al., 2011; Hoare et al., 2016). There is also some evidence that p53 is inhibiting the SASP, with trends toward increased IL1B (p value < 0.05), IL6 and IL8 in p53 inhibited cells, particularly in DDIS where p53 levels are higher (**Figure 10D-F**). However, fascinatingly when we looked at the studies where RelA (p65) had been inhibited, the results demonstrated the opposite effect. As expected, RelA inhibition showed reduced levels of IL1B and to a lesser extent IL6 (**Figure 10G-H**), suggesting reduced inflammatory signalling. However, both p53 mRNA levels and p53 activity (as represented by MDM2, p21 and GADD45A mRNA levels) were all reduced by RelA inhibition (**Figure 10I-L**). This strongly suggests that the reduced p53 signalling in OIS is not due to the increased inflammatory signalling, and makes it difficult to explain why p53 activity might peak at four days as has been suggested (Robles & Adami, 1998), before upregulation of the SASP. Reduced p53 signalling in response to RelA KD is not a phenomenon we have observed in the literature. In fact, one study by Chien et al. (2011) investigated the impact of a RelA KD in senescence and found at the protein level when RelA was knocked down, total p53 and p21 expression remained similar to that of control senescent cells.

NF-κB is a major transcriptional regulator of senescence (Oeckinghaus & Ghosh, 2009). In senescence, the NF-κB subunit RelA has been shown to be commonly accumulated on chromatin (Chien et al., 2011) suggesting the active homo/heterodimer in senescence involves RelA. As the most common subunit conformation of NF-κB is p50-RelA (Hoffmann & Baltimore, 2006), we chose to represent NF-κB as a p50-RelA heterodimer in the model network. To be considered a successful representation of cellular senescence, expression of inflammatory proteins must be higher in OIS compared to DDIS, and should not be expressed until senescence had been established at the equivalent of 5-7 days post-senescence induction (**Figure 10A-C**). Many studies additionally support the activation of the SASP as occurring between days 5-8 post senescence induction (Coppé et al., 2008; Freund et al., 2011; Hoare et al., 2016).

Inflammatory proteins were introduced into the network of sub-model B. Simulation of NF-κB is lower in OIS than in DDIS in sub-model B (**Figure 11A**), phosphorylated NF-κB (pNF-κB) and downstream target IL6 are more highly expressed in OIS than in DDIS (**Figure 11B-C**), as expected from our systematic analysis (**Figure 10A**). However, inflammatory proteins are expressed immediately upon senescence induction in sub-model B simulations which is not reflected in the literature (Coppé et al., 2008; Hoare et al., 2016) or our systematic analysis (**Figure 10A-C**). The network of sub-model B lacks a mechanism which can regulate SASP induction in senescence as it only contains the different states of NF-κB (inhibited, uninhibited and phosphorylated) and one downstream target of IL6. While NF-κB is a recognised main SASP regulator, something is required to regulate NF-kB activation. One suggested mechanism which regulates SASP induction is Notch signalling (Hoare et al., 2016) which we find evidence of at the transcript level with HES1 expression (**Figure 8C**). As previously mentioned, Notch signalling is introduced in the Senescence Induction Model network. When simulating the Senescence Induction Model under normal senescence circumstances, SASP induction begins once the Notch switch is induced at the equivalent of day 5 post senescence induction (**Figure 11D-F**), and there is a stronger inflammatory phenotype in OIS compared to DDIS as pNF-κB and IL6 expression are both higher (**Figure 11E-F**). Although NF-κB levels remain similar in OIS and DDIS, there is more pNF-κB in OIS than DDIS, which likely contributes towards the increased inflammatory response observed in OIS in comparison to DDIS, which we also see in our systematic analysis (**Figure 10A-C**).

**Figure 11.**
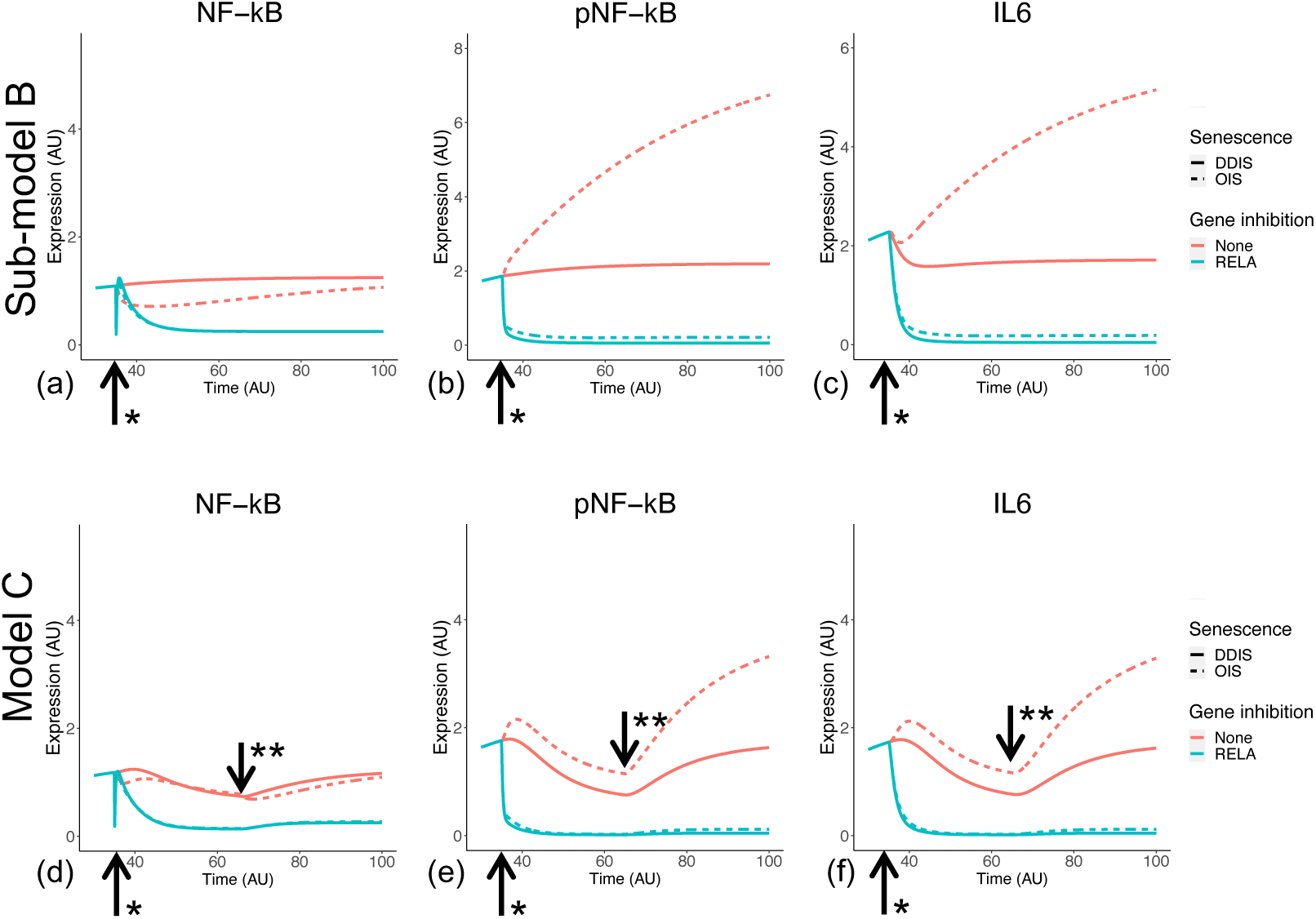
Simulation of the inflammatory SASP in cellular senescence. Simulations show temporal expression of proteins, including phosphorylation status in cases such as NF-κB, in DDIS and OIS. Units are all arbitrary. *Senescence and/or KD induced. **Notch Switch induced.

Interestingly, the systematic analysis and computational model address the issue of whether the SASP is delayed in DDIS compared to OIS, or if the secretion of SASP factors is reduced. Both support and suggest that SASP factors are not delayed in DDIS, but are rather reduced in comparison to OIS. For example, median expression of the IL6 transcript initially increases between days 5 and 7 in both DDIS and OIS, however expression has a larger range of expression in OIS than in DDIS (**Figure 10A**). The same is true at the protein level as observed in simulations of the Senescence Induction Model (**Figure 11F**). While both transcriptomic analyses and model simulations support a stronger expression of SASP proteins in OIS, it is difficult to determine whether the SASP is primarily contributing towards senescence maintenance or bystander induced senescence. However, our analyses do find persistent expression of cell cycle arrest proteins such as p53, p21 and p16.

Therefore, it could be possible that the inflammatory nature of the SASP is causing further DNA damage to sustain cell cycle arrest. Alternatively, the DNA damage initially caused by the senescence stimulus may not be resolved and could be the cause behind persistent expression of cell cycle arrest proteins.

Although expected, we could find little evidence of an initial TGFβ-rich secretome at the transcript level in DDIS or OIS. Both TGFB1 and TGFBR1 mRNAs showed no trend toward upregulation at early timepoints followed by a decrease in expression (**Figure 12A-B**). However, COL1A1, PDGFA, and ACTA2 do tend to decrease in expression from days 0-4 to days 5-7 (**Figure 12C-E**). Furthermore, the ACTA2 gene encoding the α-SMA protein, a biomarker of myofibroblast development (Wynn & Ramalingam, 2012), which is robustly expressed in response to prolonged TGFβ in both proliferating and senescent cells (Wordsworth et al., 2022), had reduced expression in OIS at 0-4 days compared to proliferating controls, while DDIS showed no change compared to proliferating cells (**Figure 12E**). If these observations are correct, the early TGFβ SASP observed by Hoare et al. (2016) may reflect normal function of proliferating fibroblasts that is reduced as Notch signalling declines and the inflammatory SASP activates.

**Figure 12.**
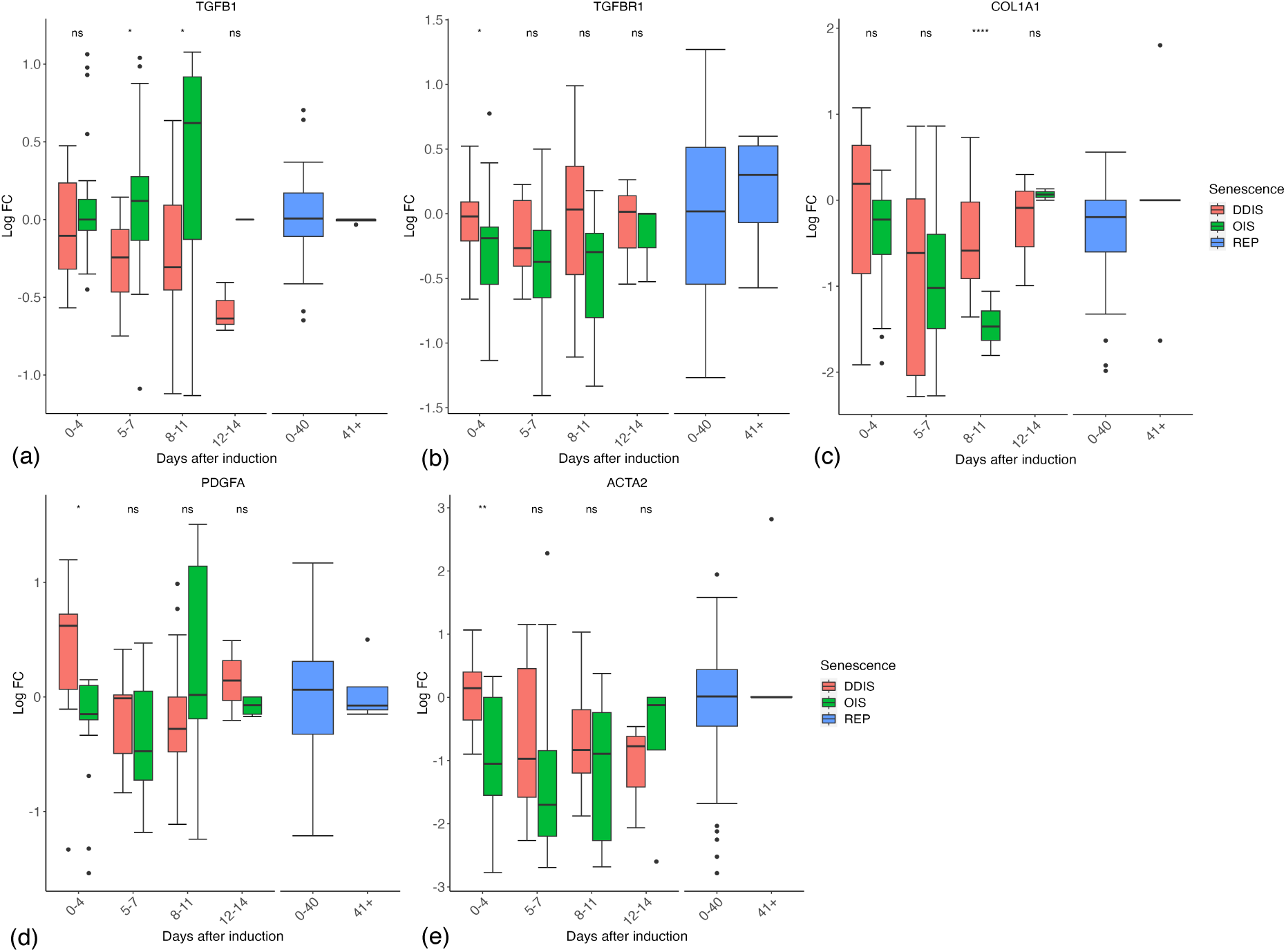
Expression of TGFβ response genes in senescent cells. (A-E) Gene expression during the timeline of senescence induction measured in days after the initial stimulus. DDIS, DNA damage induced senescence; OIS, oncogene induced senescence; REP, replicative senescence; IQR, interquartile range; LogFC, log fold change; p value refers to significance in expression between DDIS and OIS, * p value < 0.5; ** p value < 0.01; *** p value < 0.001; **** p value < 0.0001.

Although we saw no evidence of an initial TGFβ SASP in OIS or DDIS, the data are still far from conclusive, and of course only reflects the senescence profile to the extent it is determined at the transcriptional level. That said, the level of secreted proteins (as would be expected of SASP components) is perhaps well observed at the transcriptional level as the proteins may be quickly secreted and must be replaced by translation of mRNA.

We intended to include TGFβ signalling and extracellular matrix (ECM) proteins in the computational model following results from Hoare et al. (2016). However, after looking further into the literature we found minimal knowledge regarding expression of proteins such as TGFβ, SMAD3 and SMAD7 at single timepoints let alone multiple timepoints in senescence. Furthermore, the transcriptomic analysis did not find any evidence of an early fibrogenic SASP (**Figure 12**). Together this lack of protein and transcript data guided our decision to not include an early fibrogenic SASP in the Senescence Induction Model. We do not argue that there is no expression of TGFβ or related proteins in senescence, just that it is poorly understood and there is not data enough to justify the presence of an early fibrogenic SASP. Although, if more evidence were to become available, this computational model of cellular senescence can readily incorporate these dynamics. When modelling an early fibrogenic SASP within the model we were able to recapitulate the results seen in Hoare et al. (2016). From these simulations pertaining to TGFβ signalling, there were no predictions outside of the Hoare et al. (2016) study. Furthermore, the introduction of TGFβ signalling and ECM proteins into the computational model had no effect on other parts of the network. Therefore the presence of these dynamics may only represent one study rather than cellular senescence as a whole, and as previously discussed due to the minimal evidence for an early fibrogenic SASP we chose not to include this in the final model.

## Conclusion

Here, we have conducted a systematic analysis of all available transcriptomic data for senescent fibroblasts that met pre-specified inclusion criteria and qualitatively modelled temporal protein changes in senescence, including KD interventions.

A total of 12 phenotypic criteria were devised to describe normal cellular senescence and when a p53 KD or RelA KD was introduced (**Table 3**). All three model networks were tested for the capability of meeting all criteria, with the Senescence Induction Model been the only network to meet all phenotypes. Although this network is not exhaustive of all aspects of cellular senescence, the fact that the model is able to recapitulate different temporal phenotypes found in the literature and in our systematic analyses, supports the strength in the network developed. Dynamic sensitivity analysis was also performed on both DDIS and OIS simulations in the Senescence Induction Model to determine if proteins species were appropriately sensitive to parameters at different times throughout the senescence process, and it was found that most proteins are appropriately sensitive to direct inputs which induce either senescence or the activation of the Notch switch (**Figure S1**), suggesting that treatment with the same drug but at different doses may result in different outcomes in the transcriptome and therefore at the protein level. In our database, for example, there are nine studies which induce DDIS using etoposide treatment. However, there are three differing doses of etoposide treatment with different treatment regimens depending upon the study design, all which could lead to changes in the senescence phenotype. Interestingly, this modelling and sensitivity analysis, in addition to the systematic analysis, demonstrate that the temporal profile of a senescent cell is highly sensitive to the stimulus which can result in differing levels of expression.

Additional studies are still required to address how senescence changes over time, particularly at late timepoints. There were multiple variables changed between studies, which likely explains the lack of predictable biomarkers. Senescence is not a singular defined response, and the senescent phenotype depends on the context of stimulus, cell type, and timepoint among others. As demonstrated with the computational model, the different stimuli for DDIS and OIS results in differing levels of protein changes temporally (particularly in inflammatory SASP components), however there are still common trends between the two types of senescence such as expression of cell cycle inhibitors upon senescence stimulus induction and the SASP becoming active once senescence is established, also observed in the systematic analysis. In conclusion, the results of this systematic analysis suggest that while individual transcripts may not be expressed or repressed with sufficient universality to be used as universal biomarkers of senescence, they do follow predictable profiles depending on the type of senescence and time after induction, which can be modelled computationally. Furthermore, the interplay of different signalling pathways in senescence is a complex temporal process which is yet to be fully understood, with sub-model B illustrating how not including one signalling pathway can lead to incorrect simulation of protein profiles. Overall, this demonstrates the importance of looking at cells as a whole and not just individual pathways disconnected from one another.

Importantly, only data from human fibroblasts have been included in the transcriptomic database and protein behaviour in senescence simulations was guided only from studies which used human fibroblasts. This is important as although there is obvious overlap between humans and animal models such as mice and drosophila melanogaster, there are also notable differences – for example, mice have longer telomeres than humans but shorter lifespans (Calado & Dumitriu, 2013). Therefore, it is better to use human based data when wanting to ultimately understand any process in humans. The establishment of a framework for common use to identify and study senescent cells could improve the consistency between experimental analysis, which would hopefully lead to more detailed and reproducible analysis. Furthermore, we have demonstrated computationally that proteins are highly sensitive to the senescence stimuli, providing further evidence that if techniques were standardised then consistency across between experimental analysis could be improved. Here we have created a publicly available easy to use online database to allow further analysis of these 119 combined datasets and expand on the details discussed here. Furthering our understanding of the intricacies and differences in cellular senescence can only increase our chances of producing life-extending senotherapeutic technologies.

## Author Contributions

Systematic review was conducted independently by James Wordsworth and Rebekah-Louise Scanlan, RNA-seq data was collected and converted by James Wordsworth and Rebekah-Louise Scanlan, and microarray data by Louise Pease. Data was analysed by James Wordsworth and Rebekah-Louise Scanlan. Hannah O’Keefe created the online version of the database and Rebekah-Louise Scanlan fine-tuned the website. Computational modelling was performed by Rebekah-Louise Scanlan. Alvaro Martinez-Guimera performed the dynamic sensitivity analysis of the computational model. Work was developed, directed, and supervised by James Wordsworth and Daryl Shanley.

## Acknowledgements

This work was funded by NC3Rs (NC/ S001050/1) and Novo Nordisk Fonden Denmark (NNF17OC0027812).

## Data acknowledgment statement

No original data was produced or used in these analyses. Instead, all data utilised was from published studies of which the details (including accession numbers) can be found in **Table 2**.

