## Supplementary figures and tables for "Systematic transcriptomic analysis and temporal modelling of the senescent human fibroblast"

**Supplementary figures/tables**

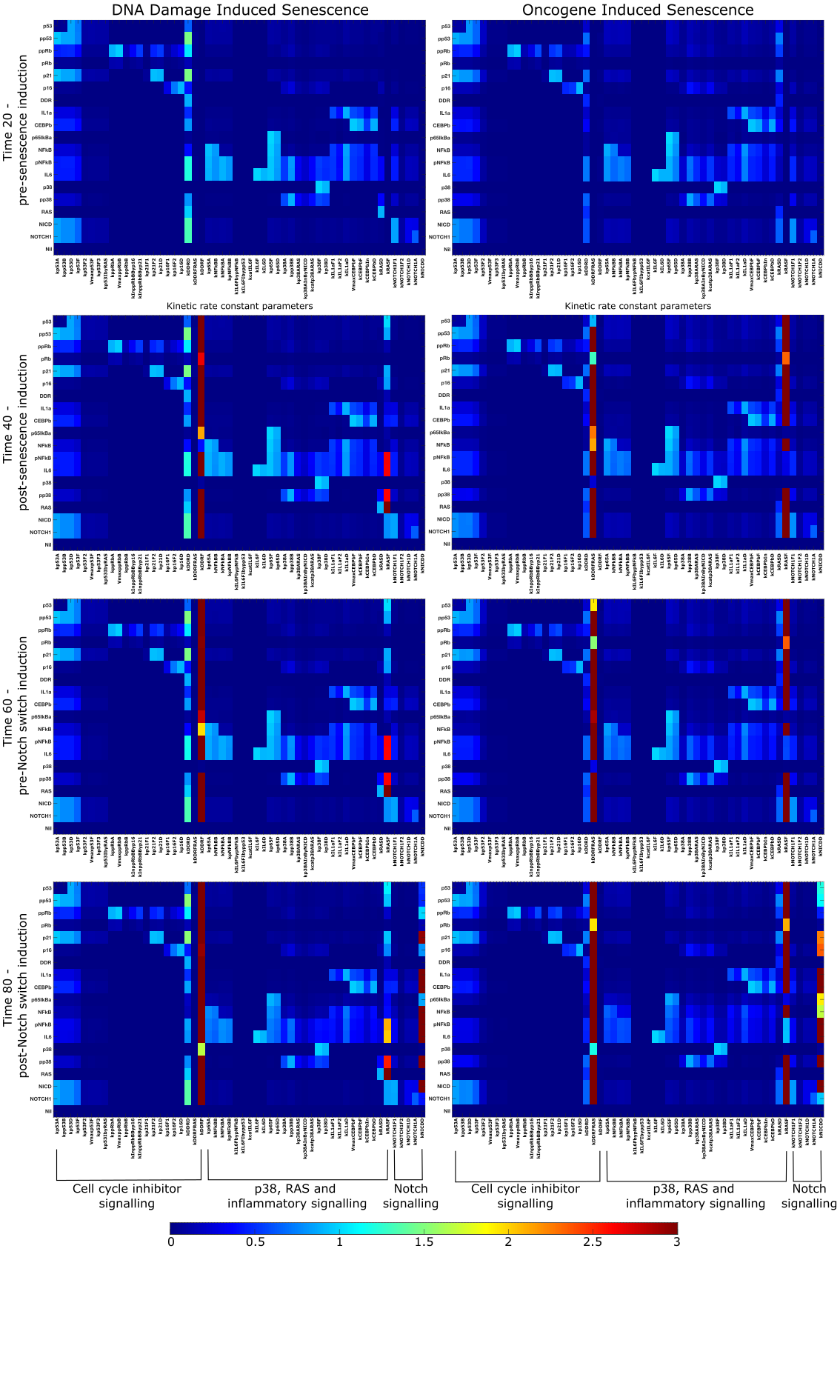

Figure S1| Dynamic sensitivity analysis of senescence simulations. Four timepoints were selected in the simulation period of DDIS and OIS, and dynamic sensitivity analysis was performed on all parameters.

Figure S2 | Dot plot of pathways from GSEA showing pathways that are activated and suppressed between (a) the 918 genes common to DDIS and OIS (p value < 0.05), (b) the 1000 genes common to DDIS and REP (p value < 1), (c) and the 137 genes specific to only BYS (p value <1). P value refers to the significance of the overrepresentation of the pathway and count reflects the number of genes associated with the pathway. DDIS, DNA damage induced senescence; OIS, oncogene induced senescence; REP, replicative senescence; BYS, bystander senescence.

Figure S3 | Gene expression of ATR, CHEK2 and CDC25A during the timeline of senescent induction measured in days after the initial stimulus. DDIS, DNA damage induced senescence; OIS, oncogene induced senescence; REP, replicative senescence; LogFC, log fold change; p value refers to significance in expression between DDIS and OIS; * p value < 0.5; ** p value < 0.01.

Figure S4 | Expression of TP53, GADD45A and GADD45B in DDIS and OIS when there is and is not p53 inhibition. Control groups for inhibition include all data for days 1-11. DDIS, DNA damage induced senescence; OIS, oncogene induced senescence; LogFC, log fold change; p value refers to significance in expression between with and without gene inhibition, ** p value < 0.01.

Figure S5 | Comparisons between DDIS and OIS across different cell lines for selected genes. We compared the gene expression for DDIS and OIS data across cell lines. All boxplots have undergone the same filtering as main text figures. DDIS, DNA damage induced senescence; OIS, oncogene induced senescence; LogFC, Log Fold Change.

| **To select for studies including fibroblasts:** | | **Search date:** | | |
| --- | --- | --- | --- | --- |
|  |  | 10 August 2020 | 06 July 2022 | 05 October 2023 |
| 1 | Fibroblasts[MeSH Terms] | 100737 | 149141 | 167132 |
| 2 | *fibroblast | 100737 | 149141 | 167132 |
| 3 | *fibroblasts | 100737 | 149141 | 167132 |
| 4 | “HCA2” OR “HCA” OR “HFF” OR “HFFF” OR “HFFF2” OR “WS1” or “Tig3” | 2892 | 5432 | 6130 |
| 5 | “BJ” OR “MRC5” OR “MRC-5” “WI-38” OR “WI38” OR “NHF” OR “NHDF” | 1381 | 1974 | 2089 |
| 6 | “IMR90” OR “IMR-90” | 8116 | 9349 | 9969 |
| 7 | #1 OR #2 OR #3 OR #4 OR #5 OR #6 | 106137 | 157313 | 175870 |
| **To select for studies looking at cellular senescence:** | | | | |
| 8 | senesce* | 5395 | 10037 | 11891 |
| 9 | senescing | 80 | 84 | 87 |
| 10 | Cellular Senescence[MeSH Terms] | 0 | 0 | 0 |
| 11 | Aging[MeSH Terms] | 11870 | 17857 | 21258 |
| 12 | Ageing | 11870 | 17857 | 21258 |
| 13 | aging | 11870 | 17857 | 21258 |
| 14 | Arrest* | 12583 | 18586 | 22443 |
| 15 | “young” AND “old” | 9845 | 15392 | 17588 |
| 16 | #8 OR #9 OR #10 OR #11 OR #12 OR #13 OR #14 OR #15 | 32195 | 51897 | 61202 |
| **To combine:** | | | | |
| 17 | #7 AND #16 | 5063 | 6175 | 7281 |

Table S1 – Systematic search terms and search results.

| Category | No.of studies | Category | No. of studies |
| --- | --- | --- | --- |
| Senescence type | | | |
| OIS | 54 | DDIS | 45 |
| REP | 24 | OSKM | 2 |
| RiboMature | 1 | BYS | 3 |
| dNTP | 1 | CR | 2 |
| NBIS | 1 | NIS | 1 |
| RNIS | 1 | MitoSkip | 1 |
| PIIPS | 1 | Trehalose | 1 |
| Control Condition | | | |
| Prolif | 111 | Quiesce | 13 |
| Immortal | 4 | Apoptosis | 1 |
| Timepoint group | | | |
| 0-4 days* | 27 | 5-7 days* | 40 |
| 8-11 days* | 25 | 12-14 days* | 8 |
| 15+ days* | 6 | 0-40 days** | 3 |
| 41+ days** | 2 |  |  |
| Genes up | | | |
| None | 118 | ZFP36L1 | 1 |
| GR | 1 | E1A | 2 |
| SmallT | 1 | Unknown | 1 |
| E6_E7 | 1 | SV40smallT | 1 |
| SmallT_E6_E7 | 1 | TGFb | 1 |
| Parkin | 3 | E7 | 1 |
| DOT1L | 1 | TRF2 | 1 |
| WSTF | 1 | Parkin_mtDNA | 1 |
| Genes down | | | |
| None | 115 | p21 | 3 |
| mTOR | 2 | pRb | 4 |
| E2F7 | 1 | pRb_E2F7 | 1 |
| p38 | 1 | ETS1 | 1 |
| JUN | 1 | RELA | 4 |
| p16 | 2 | p107 | 1 |
| p130 | 2 | H2AJ | 3 |
| mitochondrial | 2 | cGAS | 1 |
| G3BP1 | 1 | CEBPb | 1 |
| ABCD4 | 1 | AKR1C1 | 1 |
| ALOX5 | 1 | ASB15 | 1 |
| BPIL1 | 1 | BRD8 | 1 |
| C20 | 1 | CCL23 | 1 |
| CTDSPL | 1 | DCAMKL3 | 1 |
| DUSP11 | 1 | EMR4 | 1 |
| ERCC3 | 1 | GPRC5D | 1 |
| HSPC182 | 1 | IFNA17 | 1 |
| IL15 | 1 | IL17RE | 1 |
| ITCH | 1 | KCNA5 | 1 |
| KCNQ4 | 1 | LOC399818 | 1 |
| LOC51136 | 1 | MAP3K6 | 1 |
| MCFP | 1 | NRG1 | 1 |
| PEO1 | 1 | PLCB1 | 1 |
| PPP1CB | 1 | PROK2 | 1 |
| PTBP1 | 3 | PTPN14 | 1 |
| RNF6 | 1 | SHFM3 | 1 |
| SKP1A | 1 | TMEM219 | 1 |
| UBE2V21 | 1 | p53 | 7 |
| EXOC7 | 1 | LTR2 | 1 |
| LTR10 | 1 | caspase | 1 |
| IONpump | 1 | DINO | 1 |
| miR34a | 1 | unknown | 1 |
| IL1R | 1 | Complex I | 1 |
| laminA | 1 | E2F | 1 |
| HMGA1 | 1 | p130_pRb | 1 |
| p300 | 1 | CBP | 1 |
| BRD4 | 2 | p16_p21 | 1 |
| p53_pRb | 1 | ARID1B | 1 |
| HDAC | 1 | XPO7 | 1 |
| DOT1L | 1 | MIR31HG | 1 |
| YBX1 | 1 | METTL14 | 1 |
| CBS | 1 | NF1 | 1 |
| COX2 | 1 | NTRK2 | 1 |
| BDNF | 1 | COPB2 | 1 |
| ABCA1 | 1 | BAFF | 1 |
| BAK_BAX | 1 |  |  |
| Cell line | | | |
| IMR | 59 | WI38 | 19 |
| BJ | 16 | MRC | 8 |
| LF1 | 1 | HCA2 | 3 |
| FL2 | 1 | Tig3 | 3 |
| 10-5_12-1 | 1 | HDF161 | 1 |
| HFF | 5 | HMF3A | 1 |
| CAF | 3 | LFS_MDAH041 | 1 |
| HDF | 5 | Primary_fibroblast | 1 |
| HFL1 | 2 |  |  |
| Organ | | | |
| Skin | 34 | Lung | 90 |

*Table S3 | Number of studies including. *DDIS and OIS **REP only. OIS, oncogene induced senescence; DDIS, DNA damage induced senescence; REP, replicative senescence; OSKM, senescence induced as a by-product of pluripotency induction via transcription factors Oct4, Sox2, Klf4 and c-Myc; RiboMature, senescence induced through ribosomal disruption; BYS, bystander induced senescence; dNTP, depletion of deoxyribonucleotide triphosphates; CR, chromatin remodelling induced senescence; NBIS, nuclear breakdown induced senescence; NIS, Notch induced senescence; RNIS, Ras and Notch induced senescence; MitoSkip, senescence induced by skipping mitosis; PIIPS, proteasome inhibition-induced premature senescence; Trehalose, Trehalose induced senescence; Prolif, proliferating cell; Quiesce, quiescent cell; Immortal, immortalised cell; CAF, cancer associated fibroblasts.*
